# Molecular Insights into the Regulation of GNPTAB by TMEM251

**DOI:** 10.1101/2024.12.05.627003

**Authors:** Xi Yang, Balraj Doray, Varsha Venkatarangan, Benjamin C. Jennings, Danielle Henn, Jiaxuan Liang, Haikun Zhao, Weichao Zhang, Bokai Zhang, Linchen Yu, Liang Chen, Stuart Kornfeld, Ming Li

## Abstract

In vertebrates, newly synthesized lysosomal enzymes traffick to lysosomes through the mannose-6-phosphate (M6P) pathway. The Golgi membrane protein TMEM251 was recently discovered to regulate lysosome biogenesis by controlling the level of GlcNAc-1-phosphotransferase (GNPT). However, its precise function remained unclear. In this study, we demonstrate that TMEM251 is a two-transmembrane protein indispensable for GNPT stability, cleavage by Site-1-Protease (S1P), and enzymatic activity. We reconcile conflicting models by showing that TMEM251 enhances GNPT cleavage and prevents its mislocalization to lysosomes for degradation. We further establish that TMEM251 achieves this by interacting with GOLPH3 and retromer complexes to anchor the TMEM251-GNPT complex at the Golgi. Alanine mutagenesis identified F^4^XXR^7^ motif in TMEM251’s N-tail for GOLPH3 binding. Together, our findings uncover TMEM251’s multi-faceted role in stabilizing GNPT, retaining it at the Golgi, and ensuring the fidelity of the M6P pathway, thereby providing insights into its molecular function.

## Introduction

In vertebrates, most lysosomal enzymes are delivered to the organelle lumen through the mannose-6-phosphate (M6P) pathway. At the cis-Golgi, GlcNAc-1-phosphotransferase (GNPT) transfers GlcNAc-1 phosphate from UDP-GlcNAc to specific mannose residues on the high mannose glycan chains of lysosomal enzymes, following which an uncovering enzyme (UCE) removes GlcNAc, forming an M6P monoester. At the trans-Golgi, M6P receptors (M6PRs) recognize the M6P modification and selectively bind to these enzymes. M6PRs traffick from the trans-Golgi network (TGN) to the endosomes, releasing the lysosomal enzymes due to increasing acidity. These enzymes further reach the lysosome through endomembrane trafficking, while the MPRs are recycled back to the TGN through the retromer machinery^1,2^. Disruption of the M6P pathway results in the mistargeting of most lysosomal enzymes and a severe lysosome storage disease, Mucolipidosis type II (MLII)^3^.

TMEM251/GCAF/LYSET is a recently identified gene essential for the M6P pathway and the proper trafficking of lysosomal enzymes^4–7^. Patients with a loss-of-function mutation of this gene die in childhood or early adulthood. They present symptoms similar to those seen in MLII, such as severe skeletal dysplasia, coarsened facial features, short stature, and protruding abdomen^8^. Intriguingly, research shows that cells deficient in TMEM251 are refractory to several viral infections^7^. This gene is also critical for the propagation of certain tumors under nutrient-poor conditions^6^. Therefore, TMEM251 is crucial for human health and a potential drug target for treating cancers and viral infections.

TMEM251’s role within the M6P pathway is uncertain, specifically its effect on GNPT. In early September 2022, we and two other groups published simultaneously the discovery of TMEM251 and its connection to the M6P pathway. The consensus is that TMEM251 interacts with GNPT at the Golgi, which is essential for cellular GNPT activity. GNPT is an α2β2γ2 hexamer encoded by two genes, *GNPTAB* (encoding α,β subunits) and *GNPTG* (encoding γ subunit). Site-1-Protease (S1P) cleaves the α/β precursor upon its arrival at the Golgi, activating GNPT. We observed that the deletion of *TMEM251* resulted in the loss of cleaved β subunit and the accumulation of some uncleaved precursor^5^. Therefore, we proposed that TMEM251 is critical for GNPT cleavage and activation of its enzymatic activity. However, our model did not explain the reduction of total GNPT protein levels following *TMEM251* deletion. An alternative model proposed that the processing of GNPT is unaffected following TMEM251 knockout; instead, the TMEM251-GNPT interaction is necessary for retaining cleaved GNPT at the Golgi. The transmembrane helices of GNPT contain multiple hydrophilic residues that could destabilize the enzyme in the membrane. Without TMEM251, cleaved GNPT is trafficked to the lysosome and degraded by lysosomal proteases^6,7,9,10^. However, this model seems counter-intuitive since most luminal enzymes fail to reach the lysosome in *TMEM251* knockout cells, thereby abolishing their proteolytic activity. Both models account for the absence of cleaved GNPT observed following *TMEM251* knockout, yet occur through different mechanisms.

In this study, we set out to resolve this uncertainty and determine the molecular details regarding TMEM251 function. First, we confirmed that TMEM251 is essential for the M6P pathway and lysosomal digestion. TMEM251 interacts with GNPT to regulate its cleavage efficiency, enzymatic activity, and protein stability. Deleting *TMEM251* does result in the mislocalization of GNPT to the lysosome and its degradation. Through topology analysis, we determined that TMEM251 is a 2-transmembrane protein with both termini facing the cytosol. Importantly, in addition to the N-terminal cytosolic tail of GNPT interacting with COPI, the cytosolic tails of TMEM251 associate with the COPI adaptor Golph3 and retromer to maintain the TMEM251-GNPT complex at the Golgi. Thus, we propose that TMEM251 actively contributes to maintaining GNPT Golgi localization by recruiting recycling machinery, including both coatomer and retromer.

## Results

### Reevaluating the relationship between TMEM251 and GNPTαβ in the M6P pathway

Because of the controversy, we felt it is necessary to reexamine the relationship between TMEM251 and GNPTαβ carefully. Using a GNPTAB-3xHA knockin strain in HEK293T background^5^, we confirmed a loss of the cleaved β subunit upon *TMEM251* knockout, accompanied by the accumulation of a higher molecular weight band in the knockout cells (**Fig. 1A-B**). There was a ∼72% reduction in total GNPTαβ protein levels (cleaved + uncleaved). Importantly, this reduction was not due to decreased transcription, as q-PCR results indicated comparable levels of *GNPTAB* mRNA in *TMEM251* knockout cells (**Fig. 1C**). Additionally, using an antibody recognizing the α subunit, we observed a parallel abolishment of the cleaved α subunit and a reduction in total GNPTαβ levels post *TMEM251* knockout (**Fig. 1D-E**). It’s worth noting that the α subunit, after cleavage by S1P, develops more complex-type glycans and migrates similarly to uncleaved GNPTαβ^11^. Following deglycosylation by PNGseF treatment, it migrates faster than the uncleaved GNPTαβ (**Fig. 1D**, PNGaseF treated panel).

**Figure 1:**
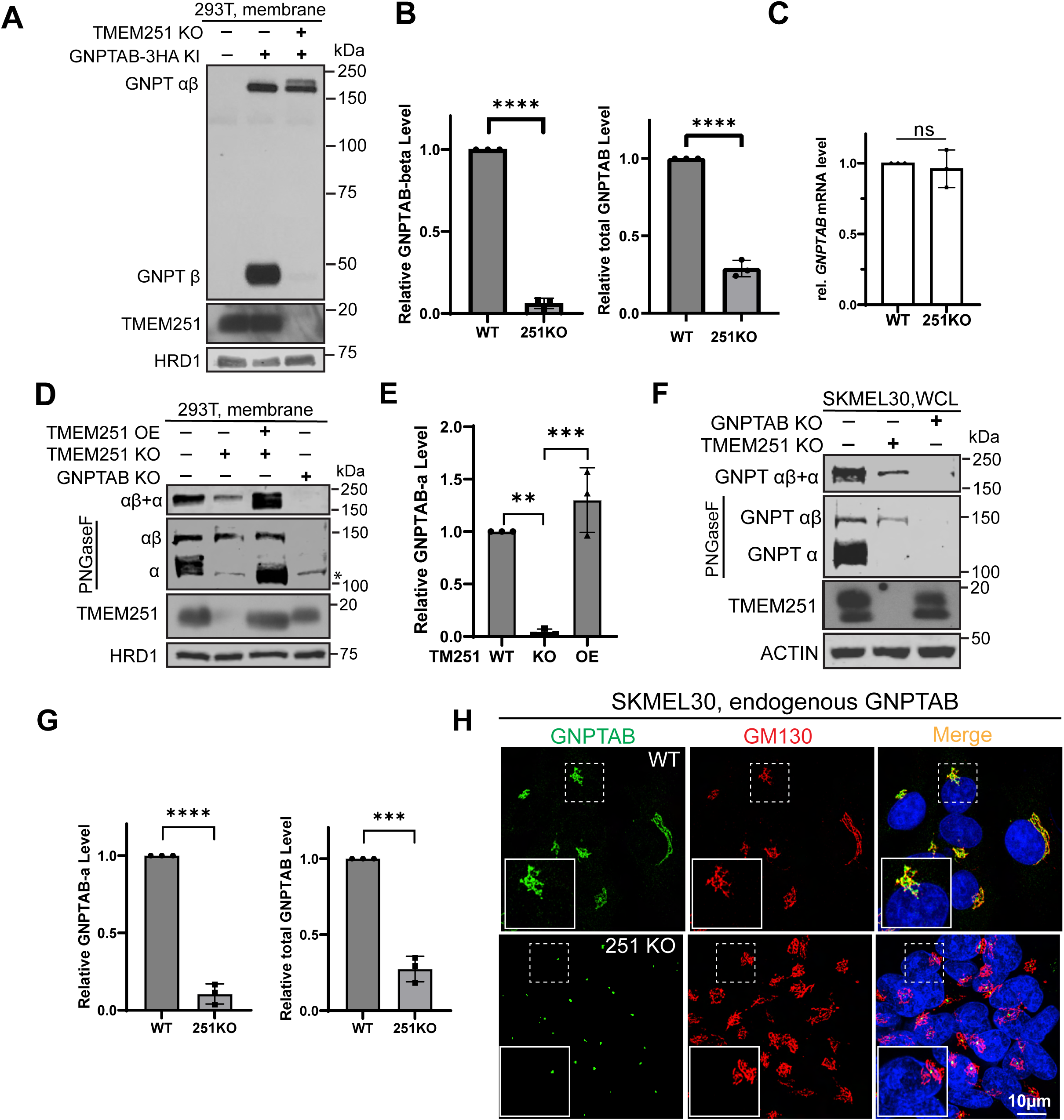
TMEM251 is critical for the stability of GNPTαβ. **(A)** Immunoblot showing the β subunit of endogenous GNPTαβ-3HA in HEK293T WT and *TMEM251* KO cells. (**B)** Quantification of (A). (**C**) qPCR analysis comparing GNPTAB mRNA levels between HEK293T WT and *TMEM251* KO cells. **(D)** Immunoblot showing the α subunit of endogenous GNPTαβ in HEK293T WT, *GNPTAB* KO, *TMEM251* KO, and TMEM251 overexpressing cells. **(E)** Quantification of (D). **(F)** Immunoblot showing the α subunit of endogenous GNPTαβ in SKMEL30 WT, *GNPTAB* KO, and *TMEM251* KO cells. WCL: whole cell lysate. **(G)** Quantification of (F). **(H)** Immunostaining images showing the localization of endogenous GNPTαβ in SKMEL30 WT and *TMEM251* KO cells.

Extending our analysis beyond HEK293T cells, we generated *TMEM251* and *GNPTAB* knockouts in the melanoma SKMEL30 cell line, known for its heightened expression of TMEM251 and GNPTαβ^7^. In line with HEK293T cell results, *TMEM251* deletion significantly reduced GNPTαβ levels (∼70% reduction) and nearly abolished the cleaved α subunit (**Fig. 1F-G**). Immunostaining using the α subunit antibody revealed that *TMEM251* deletion led to a loss of most endogenous GNPTαβ Golgi signal. However, we did not observe GNPTαβ mislocalization to lysosomes. Instead, GNPTαβ consistently shrank to a dot-like structure at Golgi, likely representing the uncleaved αβ observed in earlier immunoblots (**Fig. 1H**).

Without activated GNPTαβ, one should expect no M6P modifications on lysosomal enzymes. Indeed, using a single-chain antibody against M6P (scFv M6P)^12^, we observed no differences between samples from *TMEM251* deletion cells, *GNPTAB* deletion cells, or double deletion cells (**Fig. S1A**). The minor bands observed in knockout lanes likely represent non-specific binding by the antibody^13^. Without M6P modification, luminal enzymes, represented by CTSC and CTSD, cannot traffick to lysosomes and mature, resulting in a significant fraction being secreted into the media as unprocessed proenzymes (**Fig. S1B**). Enzyme assays of lysosomal hydrolases following CI-MPR-sepharose pull-down supported these findings, as phosphorylated β-Hex, α-galactosidase (α-Gal), and β-galactosidase (β-Gal) activity were detected only in whole cell lysates from WT cells (**Fig. S1C**). Finally, the lack of functional hydrolases was also verified by the accumulation of undigested lysosomal substrates, represented by LAPTM4A and LC3B-II (**Fig. S1D**).

In summary, using two distinct cell lines (HEK293T and SKMEL30) and antibodies targeting both α and β subunits (after 3xHA tagging), our findings consistently support that TMEM251 is essential for the processing and stability of endogenous GNPTαβ. In the absence of TMEM251, the cleaved α and β subunits are undetectable, and M6P modifications on lysosomal enzymes are absent, leading to trafficking defects of luminal enzymes and the accumulation of undigested substrates.

### The multifaceted role of TMEM251 in GNPTαβ stability, cleavage, and enzymatic activity

The absence of cleaved α and β subunits could be due to either a lack of S1P-dependent GNPTAB cleavage (processing model)^5^ or rapid trafficking of the cleaved α and β subunits to the lysosome for degradation (trafficking model)^6,7^. However, the processing model cannot explain ∼70% loss of total GNPTAB.

A key experiment to differentiate these models involves using a pulse-chase to track the fate of newly synthesized GNPTAB in the absence of TMEM251. To this end, we expressed GNPTαβ-V5 under a tetracycline-inducible (Tet-on) promoter and established stable cell lines in both WT and TMEM251 knockout backgrounds. Following a 2-hour doxycycline induction, we added cycloheximide to halt protein synthesis and monitored the fate of newly synthesized GNPTαβ-V5 at hourly intervals. In WT cells, the majority of GNPTαβ-V5 was cleaved into the β subunit, and total GNPTαβ-V5 (uncleaved + cleaved) remained relatively stable over a 5-hour chase (**Fig. 2A**). In contrast, in *TMEM251* knockout cells, significantly less β subunit was observed, but total GNPTαβ-V5 decreased by ∼60% at 5 hours (**Fig. 2A**). These results indicate that 1) the processing is nearly abolished, and 2) even uncleaved GNPTαβ-V5 is degraded in the absence of TMEM251. They are inconsistent with the hypothesis that TMEM251 has no impact on the GNPTαβ cleavage, as suggested by the trafficking model, but they do support that TMEM251 serves as an anchor to maintain GNPTαβ at the Golgi, regardless of cleavage.

**Figure 2:**
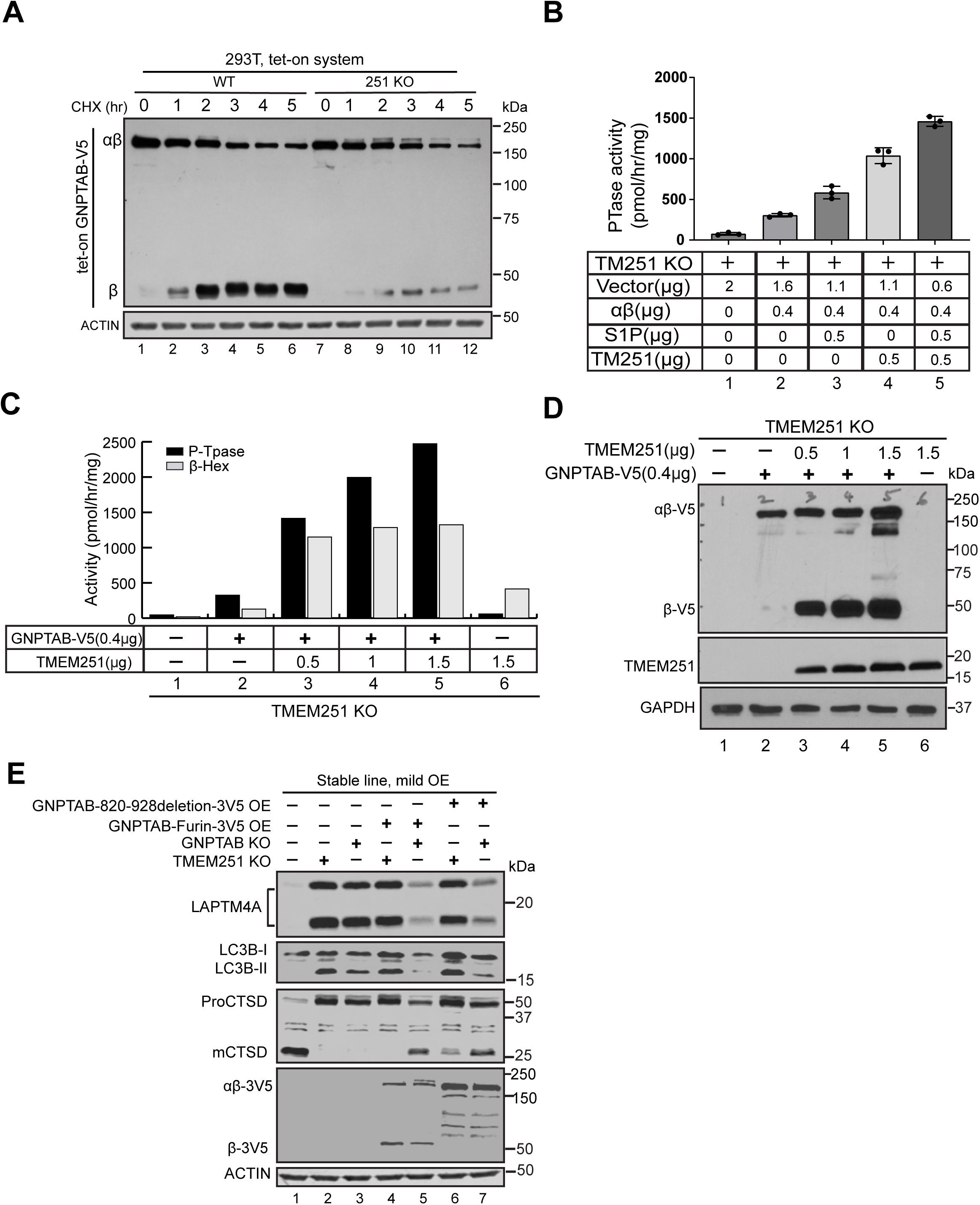
TMEM251 is essential for the processing efficiency and enzyme activity of GNPTαβ. (**A**) Representative pulse-chase experiment comparing the processing and protein stability of newly synthesized GNPTαβ in WT and *TMEM251* KO cells. The experiment was independently repeated three times. (**B**) TMEM251 and S1P work synergistically to restore the PTase activity of GNPTαβ in *TMEM251* KO cells. (**C**) Overexpression of TMEM251 in the presence of 0.4 µg of GNPTAB plasmid enhances PTase and β-Hex activities in a dose-dependent manner. Values represent the average of two assays from two independent transfections. (**D**) Under the same experimental conditions as in (C), TMEM251 overexpression increases the protein stability and processing efficiency of GNPTαβ. (**E**) Immunoblot showing that two mutants bypassing cleavage, GNPTAB-Furin-3V5 and GNPTAB(820-928)deletion-3V5, rescue *GNPTAB* KO, but not *TMEM251* KO cells.

The processing model proposes that TMEM251 acts as a bridge between S1P and GNPTαβ to enhance the cleavage of the latter. So, we asked if increasing the amount of S1P would bypass the dependence on TMEM251 to cleave GNPTαβ. To this end, we expressed GNPTαβ at low levels (400ng) in *TMEM251* knockout cells with or without overexpression of S1P and measured GNPT activity. The presence of the additional S1P clearly increased GNPT activity, nearly doubling its value (from 296 to 544 pmol/hr/mg protein, **Fig. 2B**, compare 2 & 3). However, low-level co-expression of TMEM251 with endogenous S1P had an even greater impact on GNPT activity (3.4 fold increase), whereas the greatest stimulation (almost 5 fold) occurred when both TMEM251 and S1P were co-expressed (**Fig. 2B**). These data further support that, unlike the assumption that TMEM251 has no impact on GNPTαβ cleavage, TMEM251 works synergistically with S1P to enhance its cleavage/activation efficiency.

We next asked if co-expression of TMEM251 could stabilize the GNPTαβ protein and increase its ptase activity. Keeping the amount of *GNPTAB* cDNA constant (400 ng), increasing the amount of *TMEM251* cDNA resulted in rising levels of ptase activity in a dose-dependent manner. Cellular levels of phosphorylated endogenous β-Hex also increased, but were quickly saturated (**Fig. 2C**). Further, the amount of the processed β subunit also increased in a TMEM251 dose-dependent manner (**Fig. 2D**). These results provide direct evidence that TMEM251 is critical for stabilizing GNPTαβ and enhancing its ptase activity within a cell.

Lastly, we asked if TMEM251 is required for GNPTαβ activity after its cleavage/activation. To answer this, we built a GNPTαβ construct that can bypass S1P-dependent processing by replacing its recognition sequence with that of Furin^14^. Furin is a calcium-dependent serine endoprotease that can efficiently cleave precursor proteins at the Golgi^15^, allowing efficient GNPTAB cleavage in the absence of TMEM251(Fig. 2E, lane 4 and 5). As an alternative approach to activating GNPTαβ, we directly deleted its auto-inhibition motif and S1P cleavage site (amino acids 820-928)^14^. As shown in **Figure 2E**, when stably overexpressed, both constructs can rescue the mutant phenotypes of *GNPTAB* knockout cells but not that of *TMEM251* knockout cells, indicating that TMEM251 is required for the GNPTαβ activity *in vivo* even after its cleavage. In summary, we propose that TMEM251 functions at three distinct stages to regulate GNPTαβ: 1) TMEM251 is essential for the protein stability of GNPTαβ, regardless of its cleavage state, 2) during cleavage, TMEM251 enhances the efficiency of S1P, and 3) after cleavage, TMEM251 remains crucial for the enzymatic activity of activated GNPTαβ.

### Both cleaved and uncleaved GNPTαβ undergo lysosomal degradation without TMEM251

What causes the reduction of GNPTαβ after *TMEM251* knockout? In **Figure 1H**, we cannot determine if endogenous GNPTαβ is mislocalized to the lysosome. To investigate further, we performed a longer exposure of the endogenous GNPTαβ-3xHA immunoblot and detected two prominent bands between 15 and 20 kDa in *TMEM251* KO cells (**Fig. 3A**), which likely represent degradation products of GNPTαβ-3HA (**Fig. 3B**). With the longer exposure, we also observed a minor band corresponding to the β subunit (**Fig. 3A**).

**Figure 3:**
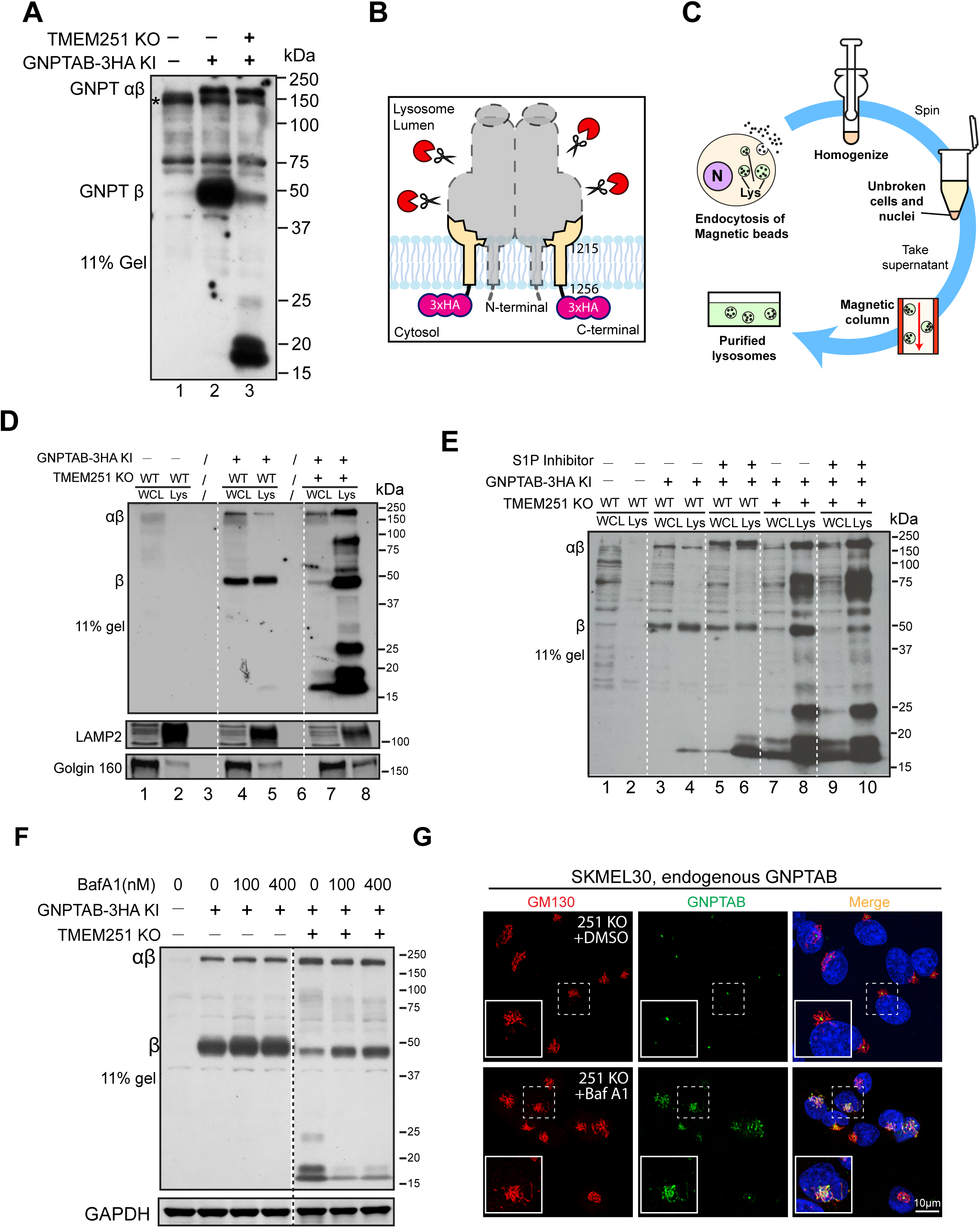
Processed and unprocessed GNPTαβ are degraded in lysosomes in *TMEM251* KO cells. **(A)** Immunoblot showing the accumulation of endogenous GNPTαβ degradation products between 15 and 20 kDa. **(B)** Model illustrating the degradation products observed in (A), likely representing the last TM helix with the 3xHA tag. **(C)** Schematic representation of magnetic isolation process of lysosomes. **(D)** Immunoblot comparing the levels of endogenous GNPTαβ-3HA in the whole cell lysate and lysosomal fractions in WT and *TMEM251* KO cells. **(E)**. Immunoblot showing the effect of S1P inhibitor on the levels of endogenous GNPTαβ-3HA in the whole cell lysate and lysosomal fractions in WT and *TMEM251* KO cells. **(F)** Protein levels of GNPTαβ in WT and *TMEM251* KO cells following BafA1 treatment. **(G)** Localization of endogenous GNPTαβ in SKMEL30 *TMEM251* KO cells after BafA1 treatment.

Since proteasomal degradation typically yields small peptides ranging from 6-10 residues^16,17^, the proteasome is unlikely to generate the observed degradation products. Therefore, we explored the possibility of lysosomal degradation. To investigate, we isolated lysosomes using magnetic beads after incubating cells with dextran-coated magnetite (DexoMAG®), which is endocytosed and delivered to the lysosome (**Fig. 3C**). In lysosomes from

WT cells, β subunit was clearly detected, while levels of the uncleaved GNPTαβ precursor were negligible (**Fig. 3D**, lanes 4 vs. 5). Conversely, lysosomes from TMEM251 KO cells contained dramatically increased amounts of both cleaved and uncleaved GNPTαβ-3xHA, accompanied by several prominent degradation intermediates (**Fig. 3D**, lanes 7 vs. 8). To answer if S1P cleavage is required for the mislocalization of GNPTαβ-3xHA, we treated cells with S1P inhibitor, PF-429242, for 19 hours before isolating lysosomes. As shown in **Figure 4E**, even though the cleaved β subunit is much reduced after PF-429242 treatment, uncleaved GNPTαβ-3xHA still was mislocalized to the lysosome, and the total level of lysosomal GNPTαβ-3xHA was largely unchanged (lanes 8 vs. 10). These results support that endogenous GNPTαβ-3xHA is mislocalized to the lysosome without TMEM251, regardless of its cleavage state.

**Figure 4:**
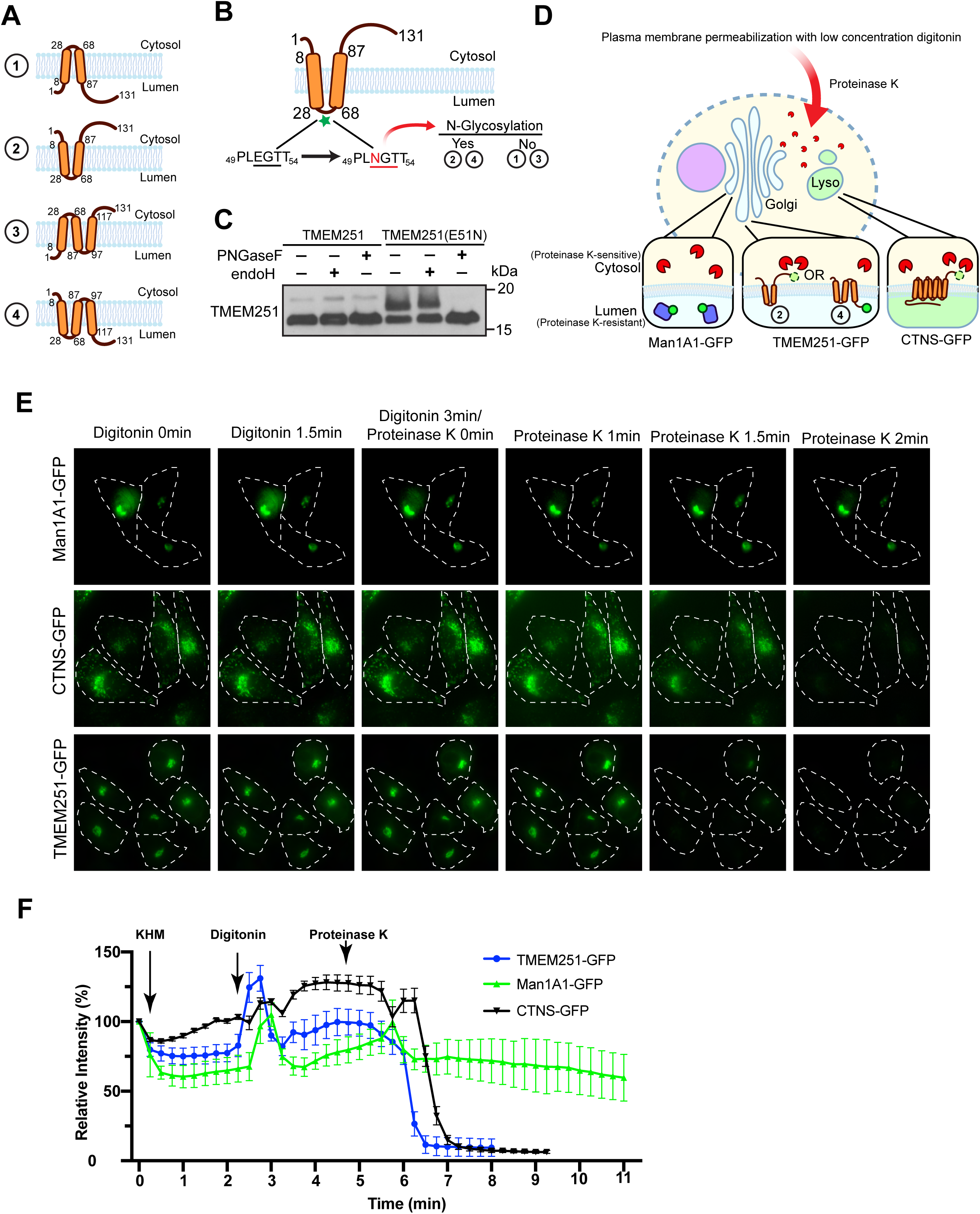
TMEM251 is a two-transmembrane helix protein with both termini in the cytosol. (**A**) Schematic representation of the four possible membrane topologies of TMEM251. (**B**) Cartoon illustrating the E51N mutation. Glycosylation of this mutant would support either topology 2 or 4 shown in (A). (**C**) Immunoblot showing the expression of TMEM251 and TMEM251(E51N) before and after treatment with either endoH or PNGaseF. (**D**) Model depicting the limited proteinase K assay. Man1A1-GFP serves as the luminal control, while CTNS-GFP acts as the cytosolic control. (**E**) The limited proteinase K assay of cells expressing Man1A1-GFP (top panel), CTNS-GFP (middle panel), and TMEM251-GFP (bottom panel). (**F**) Quantification of GFP signal from (E).

We also treated cells with BafA1 as an alternative method to confirm lysosomal degradation. As shown in **Figure 3F**, 16 hours of BafA1 treatment significantly reduced the prominent degradation products between 15 and 20 kDa and partially stabilized the β subunit. We then investigated whether BafA1 treatment would reveal lysosomal mislocalization of GNPTαβ. Surprisingly, immunostaining with the α antibody showed that the stabilized GNPTαβ was localized to the Golgi rather than the lysosome (**Fig. 3G**), suggesting that BafA1 stabilizes GNPTαβ by preventing its exit from the Golgi rather than by inhibiting lysosomal degradation. This result contrasts with previous observations that BafA1 stabilizes mislocalized GNPTαβ at the lysosome, as reported by Richards et al^7^., yet it aligns with findings in an accompanying study^6^.

In conclusion, our findings independently verified that *TMEM251* knockout leads to the mislocalization and degradation of GNPTαβ in the lysosome, regardless of its cleavage state. Interestingly, BafA1 treatment prevents GNPTαβ exit from the Golgi rather than solely blocking lysosomal degradation.

### TMEM251 has two transmembrane helices with both termini facing the cytosol

In order to understand how TMEM251 retains GNPTαβ in the Golgi, we first need to establish TMEM251’s membrane topology. AlphaFold predicted TMEM251 to be a three-transmembrane (TM) helices-containing protein^18^, while other prediction tools, such as TOPCONS and OCTOPUS, suggested it had 2 TM helices^19,20^. A recent publication proposed that TMEM251 has two TMD with both N- and C-termini facing the lumen^8^. However, no experimental evidence was presented.

Four reasonable topology models exist for TMEM251, as depicted in **Figure 4A**. We first determined if the loop between TM helices 1 and 2 is located in the cytoplasm, as was previously proposed^8^. A native ^51^EGT^53^ sequence within the loop was mutated to create an NxS/T *N*-linked glycosylation site (E51N, **Fig. 4B**) with no detrimental effect on the function of TMEM251 in stimulating GNPTαβ activity(**Fig. S2A**). The E51N mutant migrated slower than WT TMEM251, suggesting it got glycosylated. To confirm this, cell lysate was treated with either Endo H, which had minimal effect, or PNGase F, which collapsed the slower-migrating band to the size of WT, confirming that the E51N mutant had acquired complex N-linked glycans (**Fig. 4C**). This result demonstrates that the loop region located between residues 28 and 68 is in the lumen of the Golgi, ruling out topology models 1 and 3.

The C-terminal tail faces the cytosol in model 2, while in model 4, it faces the lumen (**Fig 4A**). To distinguish between these possibilities, we employed a proteinase K sensitivity assay in which cells partially permeabilized with digitonin were treated with proteinase K, which can then only digest cytosolic proteins, but not those in organellular lumens (**Fig. 4D**).

For this assay, the C-terminus of TMEM251 was tagged with GFP, which does not disrupt the normal function of TMEM251 (**Fig. S2B-C**). As controls, we used Man1A1-GFP, a Golgi lumenal protein (proteinase K resistant), and CTNS-GFP, a lysosome membrane protein with the C-terminus facing the cytosol (proteinase K sensitive). Over a 2-minute time period, Man1A1-GFP remained fluorescent, while both CTNS-GFP and TMEM251-GFP were quenched (**Fig. 4E-F**). This result shows that the C-terminal tail of TMEM251 is located in the cytoplasm, eliminating model 4. Therefore, TMEM251 contains two transmembrane helices, with both its N and C termini facing the cytosol (i.e., topology model 2).

### Alanine scan of TMEM251 reveals residues critical for its function and localization

Next, we performed a comprehensive alanine scanning analysis to identify critical residues for TMEM251 function. We systematically mutated each amino acid of TMEM251 to alanine, grouping five residues at a time. Because the anti-TMEM251 antibody only recognizes the C-terminal tail, mutations closer to the C-terminus (residues 102 to 131) cannot be detected by the TMEM251 antibody. To address this, either an N-terminal 3xFLAG tag or 3xHA tag was incorporated for immunoblotting and immunostaining.

In total, 31 mutants covering the entire more common, short-isoform of TMEM251 were expressed in *TMEM251* KO HEK293T cells under the tet-on promoter. This promoter, when uninduced, maintains a low level of protein expression (leaky expression), preventing potential overexpression artifacts. To assess the functionality of TMEM251 mutants, we utilized mature CTSD levels as a readout (**Fig. 5A**). The deletion of TMEM251 resulted in the abolishment of mCTSD (**Fig. 5A**, compare lanes 1 to 2). Reintroducing TMEM251 mutants restored CTSD maturation to varying degrees. A disease mutant, R7W, served as a positive control in this screen^8^. A 50% or less cutoff of WT mCTSD levels identified ten defective mutants with loss of TMEM251 function (highlighted in red or yellow, bottom panel, **Fig. 5A**).

**Figure 5:**
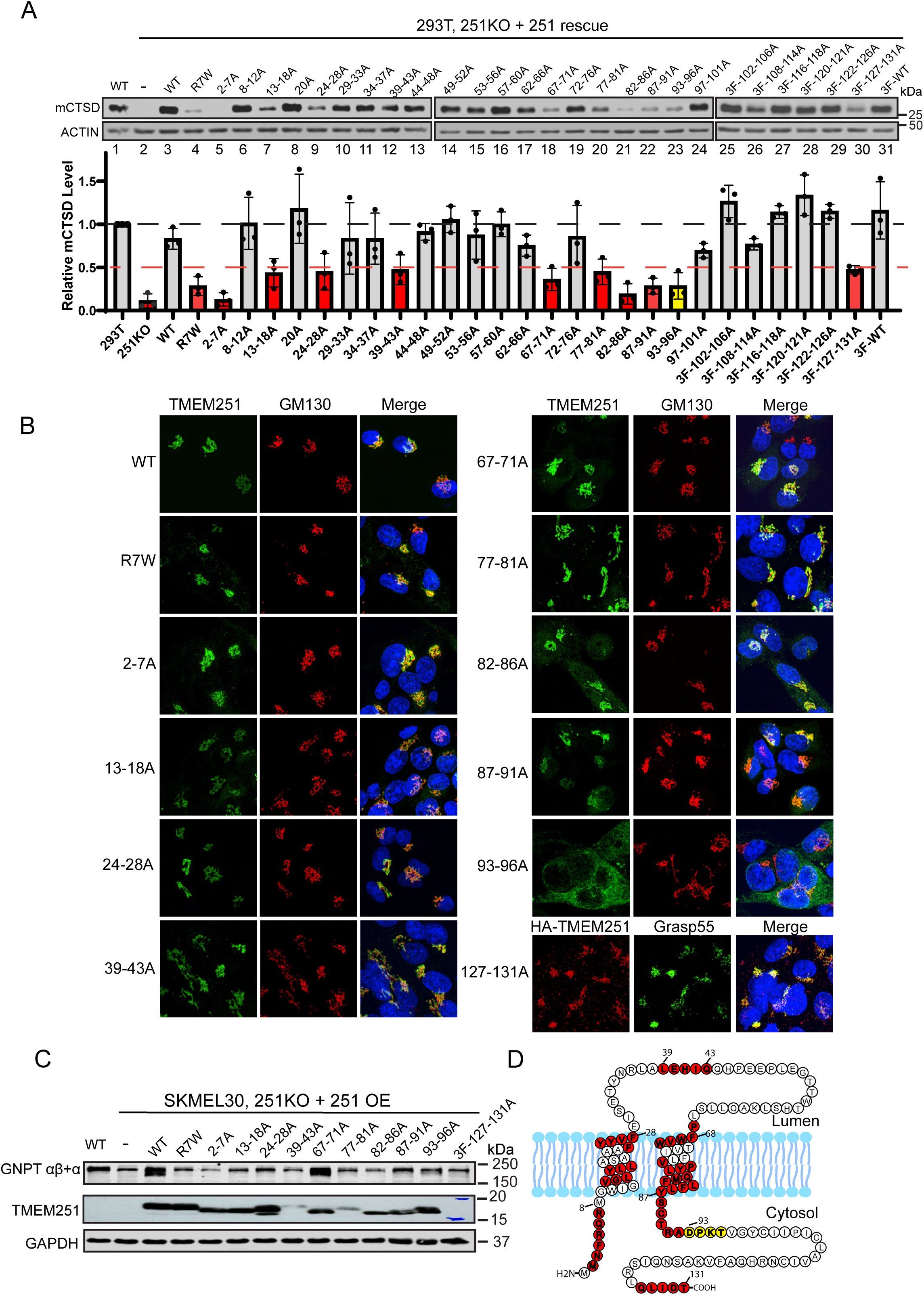
Alanine scan of TMEM251 reveals residues critical for its function. (**A**) Immunoblot analysis of mCTSD levels in HEK293T *TMEM251* KO cells complemented with various TMEM251 mutants. Mutants with mCTSD levels below 50% are marked in red, indicating a loss of function for TMEM251. The 93-96A mutant is highlighted in yellow, as discussed further below. (**B**) Immunofluorescence images showing the localization of defective TMEM251 mutants identified in (A) within SKMEL30 *TMEM251* KO cells. (**C**) Immunoblot showing the protein levels of GNPTαβ in cells expressing different TMEM251 mutants. (**D**) Topology map of TMEM251, highlighting residues critical for its function and localization in color.

The subcellular localization of these mutants was determined by stably expressing them in SKMEL30 *TMEM251* knockout cells and immunostaining. Nine out of ten mutants localized to the Golgi and co-localized with GM130 (**Fig. 5B, S3A**). Only TMEM251(93-96A) lost its Golgi localization and was trapped in the ER (**Fig. 5B and S3B**).

We also examined whether these mutations led to the loss of GNPTαβ. While all other mutations led to a significant reduction in GNPTαβ level, TMEM251(67-71A) rescued it to the same extent as WT (**Fig. 5C**). This suggests that mutations in this region do not disrupt TMEM251’s function of anchoring GNPTαβ in the Golgi, but potentially disrupted some other function of this protein in the M6P pathway.

In summarizing these results, we plotted the critical regions in the topology model (**Fig. 5D**). It indicates that the two TM helices (13-18, 24-28, 67-71, 77-81, 82-86) and the immediate cytosolic regions next to them (2-7, 87-91, 93-96) are crucial for TMEM251 function. Among them, residues 93-96 are particularly vital for the ER exit. Lastly, a small region within the lumenal loop (39-43) and the very C-terminus (127-131) also play a critical role in TMEM251 function.

### The N-terminal tail of TMEM251 interacts with Golph3 and facilitates its Golgi localization

We next sought to dissect the molecular mechanism that holds the TMEM251-GNPTαβ complex in the Golgi. The N-terminal cytoplasmic tail (N-tail) of GNPTαβ is known to bind directly to the β/δ and γ/ζ subunits of COPI, and this interaction is critical for the Golgi localization of GNPTαβ^21^. Patient mutations disrupting this interaction result in the enzyme’s mislocalization to the lysosome, leading to Mucolipidosis III^21^. However, the mislocalization of GNPTαβ observed upon TMEM251 depletion implies that the GNPTαβ-COPI interaction alone is insufficient to retain the enzyme at the cis-Golgi, underscoring the additional contribution of TMEM251. Then, how does TMEM251 perform this role?

We first determined if the N-terminal cytoplasmic tail of TMEM251 was sufficient to redirect a GFP reporter from the plasma membrane to the Golgi. For this assay, we utilized a construct, SI^N^-SI^TM^-GFP, encoding the N-tail and transmembrane segment of the plasma membrane protein, sucrase isomaltase, fused to GFP. We then switched its N-tail with the first seven amino acids of the short isoform of TMEM251 to generate 251^N^-SI^TM^-GFP (**Fig. 6A**). We and others have shown that while SI^N^-SI^TM^-GFP localizes to the plasma membrane, replacement of its N-tail with the N-tails from several other Golgi glycosyltransferases, including GNPTαβ, restricts the chimera to the Golgi^21,22^. Immunofluorescence microscopy of WT HeLa cells transfected with 251^N^-SI^TM^-GFP shows tight colocalization with the Golgi marker, giantin, whereas SI^N^-SI^TM^-GFP was predominantly on the plasma membrane (**Fig. 6B-C**). This result shows that the N-tail of TMEM251 has sufficient information to retain the chimera in the Golgi. Further alanine mutagenesis scan revealed that residues F4 and R7 are critical for the Golgi localization (**Fig. 6D-F**). Importantly, the R7W mutation in humans causes Mucolipidosis type V^5,8^.

**Figure 6:**
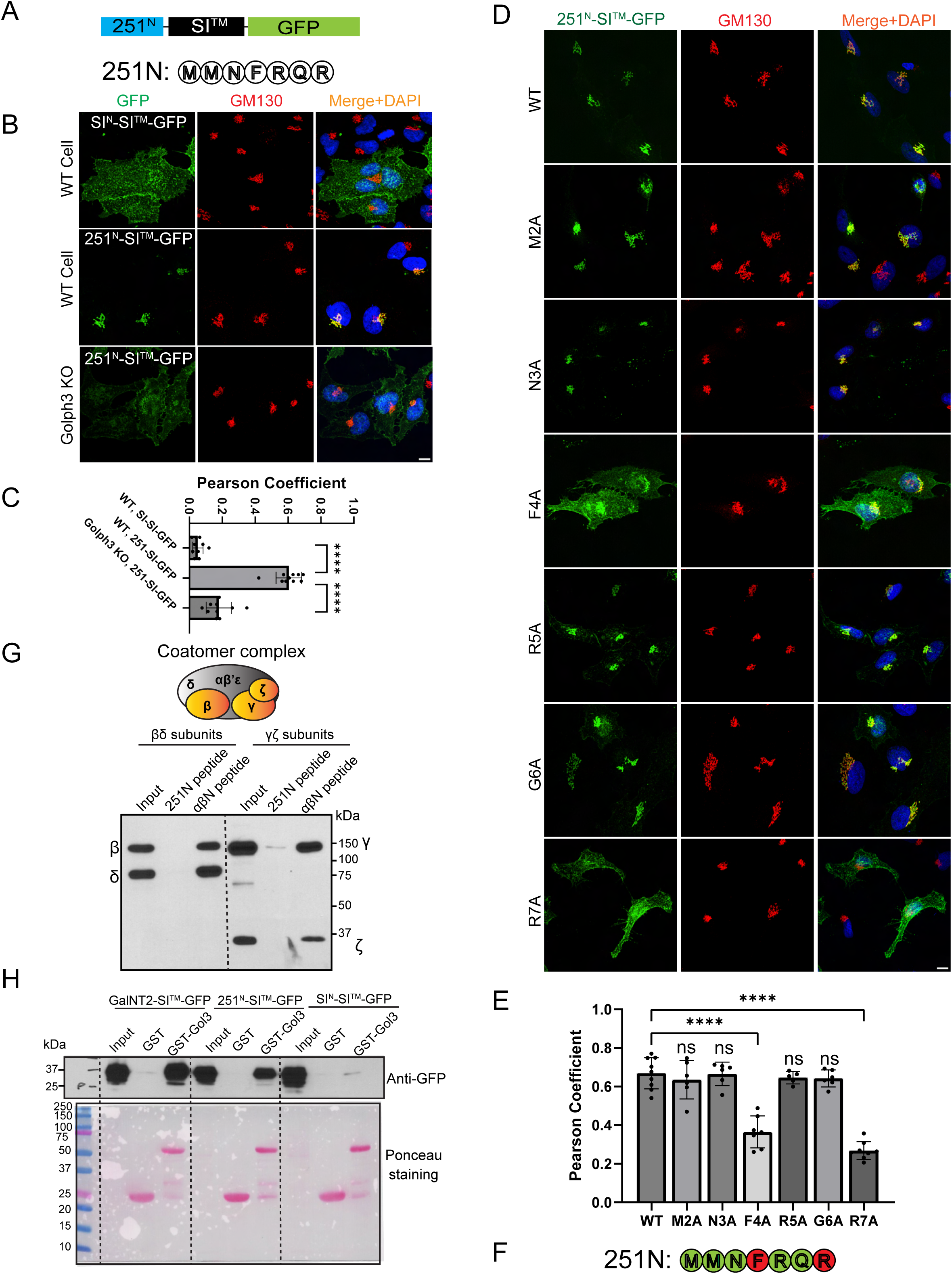
The N-terminal F^4^XXR^7^ motif of TMEM251 contributes to its Golgi localization by interacting with Golph3. **(A)** Schematic representation of the chimera. The N terminal tail of TMEM251 is fused with the transmembrane helix of sucrase isomaltase (SI), followed by GFP. **(B)** Colocalization of SI^N^-SI^TM^-GFP and 251^N^-SI^TM^-GFP with GM130 in HeLa WT and *GOLPH3* KO cells. **(C)** Pearson coefficient analysis of colocalization data from (B). (**D**) Alanine mutagenesis scan to identify critical residues required for Golgi localization. (**E**) Pearson coefficient analysis of colocalization data from (D). (**F**) The F^4^XXR^7^ motif is critical for the Golgi localization of TMEM251. (**G**) Pull-down assays using synthesized TMEM251 N-terminal peptide showed no interaction with the COPI β/δ or γ/ζ subcomplexes. The N-terminal peptide of GNPTαβ (αβN peptide) was used as a positive control. **(H)** Pull-down assays demonstrating the interaction between the TMEM251-N chimera and GST-Golph3. GalNT2-SITM-GFP and SIN-SITM-GFP served as positive and negative controls, respectively. Scale bar:10 µm.

The imaging data raised the question of which recycling machinery retains 251N-SITM-GFP in the Golgi. Given that GNPTαβ binds COPI, we hypothesized that COPI might also mediate TMEM251 retention. To test this, we synthesized a biotinylated peptide corresponding to the first 7 amino acids (N-tail) of TMEM251, immobilized it on streptavidin beads, and performed pull-down assays with Sf9 insect cell lysates expressing the β/δ or γ/ζ subcomplexes of COPI^21^. While the GNPTαβ N-tail (positive control) robustly bound both β/δ and γ/ζ dimers, no binding was detected with the TMEM251 N-tail (**Fig. 6G**).

The F^4^XXR^7^ motif in TMEM251’s N-tail suggested that its interaction with COPI might be indirect, mediated by the adaptor protein Golph3, given its similarity to the LXXR Golph3 recognition motif ^22^ ^23–25^. The Munro group has shown that a GST-Golph3 fusion protein efficiently bound GalNT2-SI^TM^-GFP that carries the N-tail of the Golgi enzyme *N*-acetylgalactosaminyltransferase2 ^22^. We prepared GST-Golph3 and performed the binding assays using GalNT2-SI^TM^-GFP and SI^N^-SI^TM^-GFP as positive and negative controls, respectively. As shown in **Fig 6H**, 251^N^-SI^TM^-GFP bound well to GST-Golph3, while there was no binding of GST-Golph3 to SI^N^-SI^TM^-GFP as previously reported ^22^.

To confirm Golph3’s role *in vivo*, we deleted *GOLPH3* in HeLa cells, which caused 251N-SITM-GFP to lose most of its Golgi localization (**Fig. 6B-C**). Knocking down GOLPH3 also reduced endogenous GNPTαβ levels and disrupted lysosomal enzyme trafficking, as evidenced by impaired CTSC and CTSD maturation (**Fig. S4A-B**). Immunostaining of SKMEL30 cells further confirmed diminished Golgi signals for GNPTαβ in GOLPH3-deficient cells (**Fig. S4C-D**).

In summary, our findings demonstrate that TMEM251’s N-tail mediates Golgi localization through Golph3 interaction. Together with our previous findings on GNPTαβ^21^, this highlights the importance of N-tail recycling for the efficient activity and retention of both proteins in the Golgi.

### The retromer complex recycles TMEM251 from the lysosome to the Golgi

Another critical molecular machinery for maintaining membrane proteins at the Golgi is the retromer complex^26–29^. Retromers actively retrieve cargo proteins, such as CI-MPR and sortilin, from endosomes and transport them back to the trans-Golgi network, preventing their degradation in the lysosome. To assess whether the retromer complex also contributes to the Golgi localization of TMEM251, we deleted *VPS35*, a key component of the VPS26/29/35 retromer subcomplex, in HEK293T cells. Deletion of *VPS35* resulted in a significant reduction in endogenous GNPTαβ protein levels, while TMEM251 levels remained largely unchanged (**Fig. 7A**). Lysosome purification from two independent knockout cell lines revealed that significant fractions of GNPTαβ and TMEM251 were mislocalized to the lysosome and partially degraded (**Fig. 7B**, lanes 6, 8 vs. 4). Furthermore, we observed a decrease in VPS26, another essential component of the retromer complex, indicating that the stability of retromer subunits is dependent on an intact retromer subcomplex (**Fig. 7A**). Similar observations have been reported before^30^.

**Figure 7:**
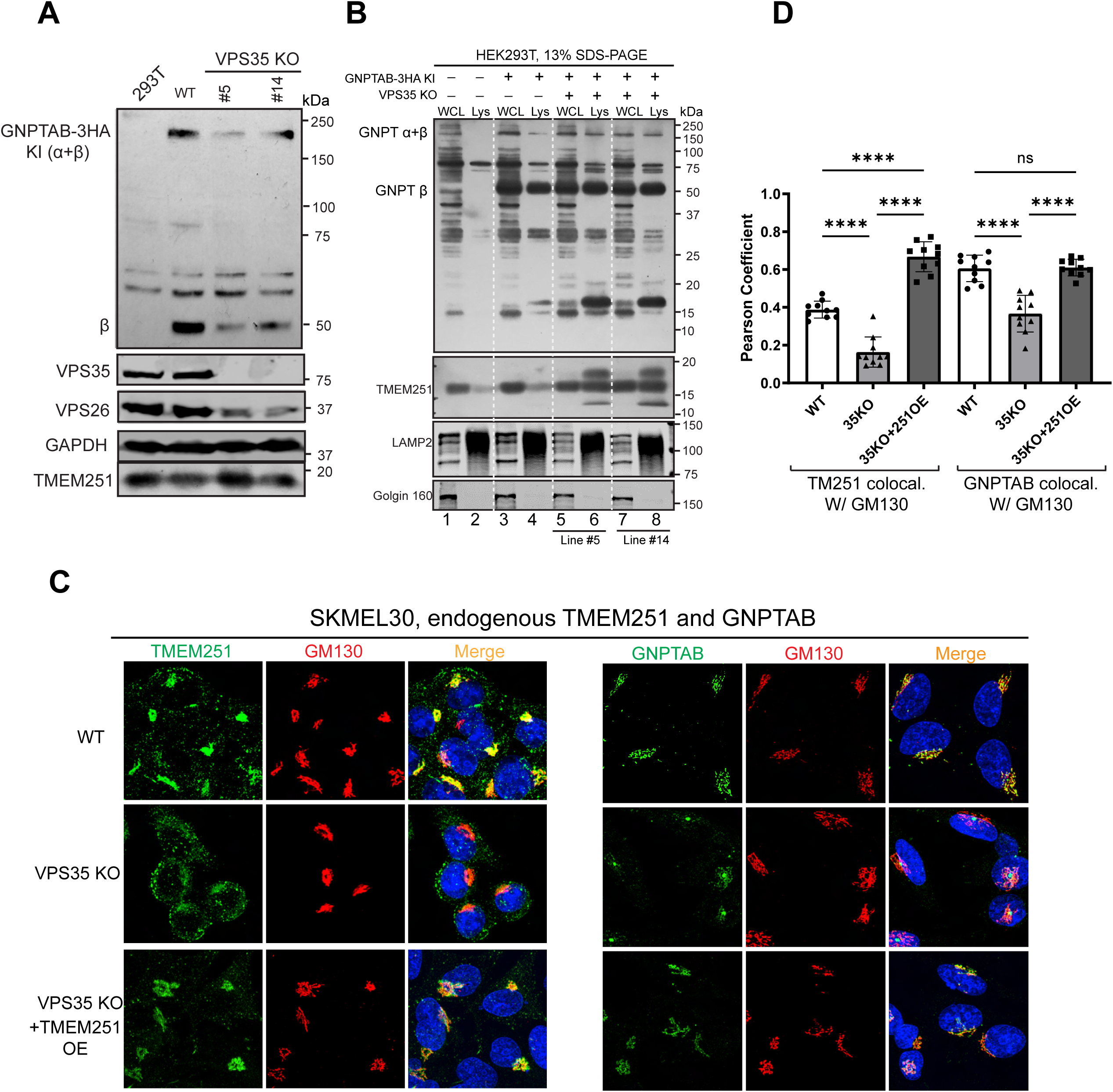
The retromer complex recycles TMEM251 from endosome to Golgi. (**A**) Immunoblot showing endogenous GNPTαβ protein levels in two colonies of *VPS35* KO HEK293T cells. (**B**) Lysosome isolation demonstrating that a fraction of GNPTαβ and TMEM251 is mislocalized to the lysosome and degraded in *VPS35* KO cells. (**C**) Localization of endogenous TMEM251 and GNPTαβ in SKMEL30 WT, *VPS35* KO, and *VPS35* KO+*TMEM251* overexpression cells. (**D**) Pearson coefficient analysis of colocalization data from (C).

Utilizing the SKMEL30 cell line, we investigated the localization of endogenous TMEM251 and GNPTαβ through immunostaining. Deletion of *VPS35* led to the mislocalization of TMEM251 from the Golgi to punctate structures (**Fig. 7C-D**). Similarly, the Golgi localization of GNPTαβ was markedly reduced (**Fig. 7C-D**), and gave the same central Golgi, single dot per cell pattern as seen for *TMEM251* KO (**Fig. 1H**). These results underscore the critical role of the retromer complex in maintaining both proteins at the Golgi.

We then explored whether overexpression of TMEM251 could restore the Golgi localization of GNPTαβ in *VPS35* deletion cells. Indeed, upon overexpression, a substantial amount of TMEM251 localized to the Golgi, leading to the restoration of GNPTαβ Golgi localization (**Fig. 7C-D**). These findings suggest that the loss of GNPTαβ Golgi localization following *VPS35* deletion is attributable to the depletion of TMEM251 from the Golgi.

Combining the results from alanine scanning, Golph3, and retromer machinery, we propose a model wherein TMEM251 actively participates in maintaining GNPT at the Golgi and regulating its cleavage and enzymatic activity (**Fig. 8**). Firstly, TMEM251 forms a stable complex with GNPT. Secondly, the N-termini of both TMEM251 and GNPT interact with the COPI machinery either indirectly (through Golph3) or directly, facilitating the efficient recycling of the TMEM251-GNPT complex from the trans-Golgi network to the cis-Golgi. Thirdly, under normal conditions, a small fraction of the TMEM251-GNPT complex may exit the Golgi and mislocalize to endo-lysosomes, but this population is efficiently recycled back to the trans-Golgi network by the retromer machinery. Thus, TMEM251 actively contributes to maintaining Golgi localization by engaging both the COPI and retromer machinery.

**Figure 8:**
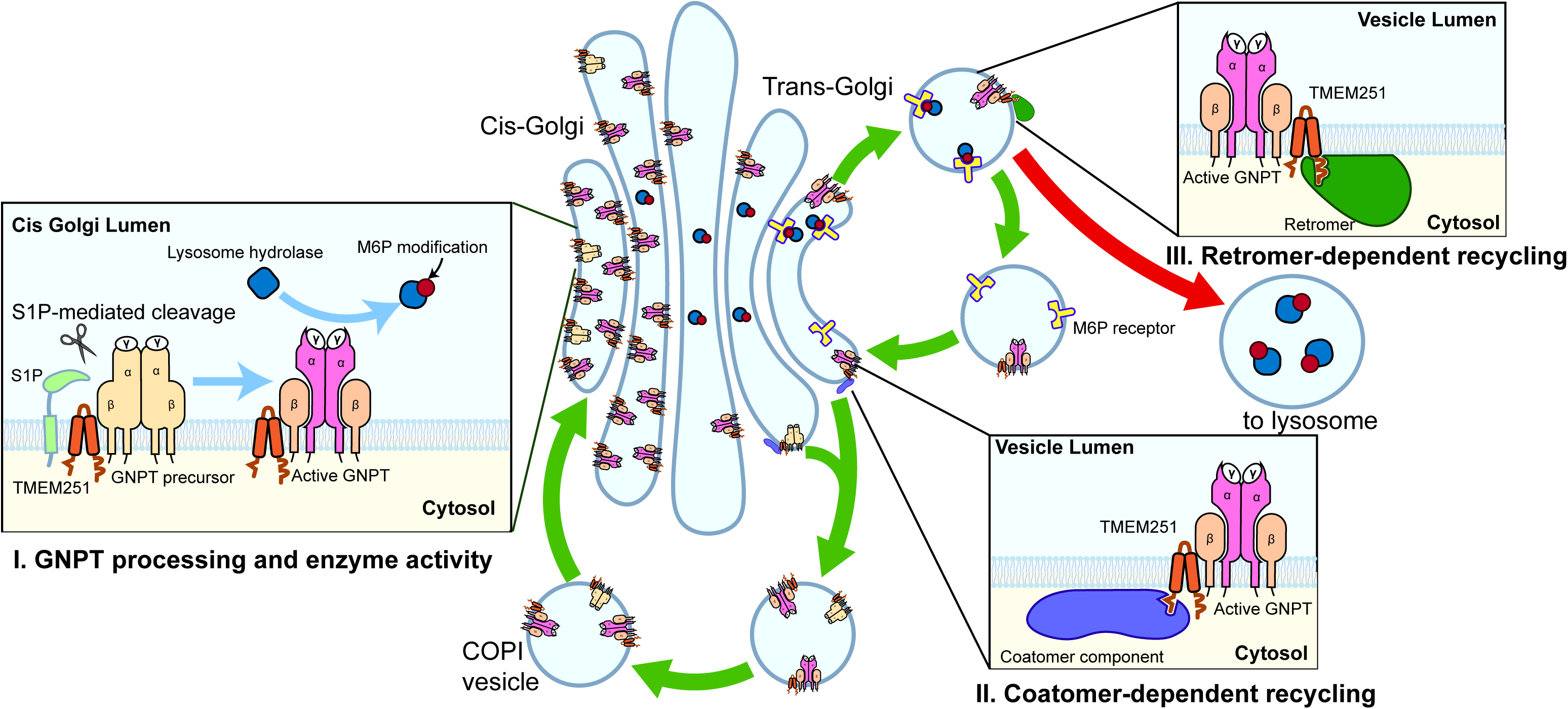
A model summarizing the role of TMEM251 in the processing, enzymatic activity, and localization of GNPT.

## Discussion

### TMEM251 is critical for GNPT activity at the physiological level

For many years, the M6P pathway was considered a well-understood mechanism for lysosome biogenesis. The study uncovering the role of COPI in the Golgi recycling of GNPTαβ and the cryo-EM work elucidating its structure seemed to close a final chapter in the field^14,21,31,32^. However, the recent discovery of TMEM251 and its indispensable role in the M6P pathway has reignited interest in this field of exploration. The phenotypes of TMEM251 ablation closely mimic the loss of GNPT, resulting in mucolipidosis type V. While there is consensus that TMEM251 plays a critical role in the M6P pathway, the specific mechanism of this regulation has been a subject of debate. In this study, we revisited the intricate details of how TMEM251 modulates the M6P pathway.

Our findings highlight the significance of TMEM251 in facilitating the efficient cleavage of newly synthesized GNPTαβ (**Fig. 2A**). Even following cleavage, TMEM251 remains crucial for its activity when GNPT is expressed at endogenous levels or mildly overexpressed (**Fig. 2E**).

This regulation might operate on two levels. Firstly, TMEM251 may act as a mediator between S1P and GNPT to enhance cleavage efficiency. In support of this model, we previously observed that TMEM251 can pull down both GNPT and S1P^5^. Without TMEM251, cleavage and activation become inefficient. Secondly, the TMEM251-GNPT interaction retains GNPT within the Golgi apparatus. In the absence of TMEM251, the uncleaved GNPT rapidly exits the Golgi, reducing the precursor concentration and diminishing S1P cleavage efficiency. However, overexpression of GNPT or S1P could increase the likelihood of interaction between GNPT and S1P, thereby bypassing the need for TMEM251 for cleavage.

### TMEM251 preserves GNPT at the Golgi

Our investigation reaffirmed the vital role of TMEM251 in maintaining GNPT at the cis-Golgi, consistent with the initial suggestions by Richards et al. and Pechincha et al.^6,7^. Using magnetic beads, we purified lysosomes and observed that in the presence of TMEM251, approximately 10% of the activated GNPT was already localized to the lysosome in WT cells(**Fig. 3D-E**). In contrast, minimal uncleaved GNPT reached the lysosome. Upon *TMEM251* deletion, a significant increase of both uncleaved and cleaved GNPT in the lysosome was observed. Notably, the majority of these mislocalized enzymes underwent proteolytic cleavage, resulting in smaller degradation products. These results also indicate that the lysosomes still retain partial proteolytic function, implying that there are other M6P-independent pathways to traffick lysosomal enzymes. Our attempts to detect GNPTαβ mislocalization through immunostaining were unsuccessful. While endogenous GNPTαβ was readily detected in WT SKMEL30 cells, *TMEM251* deletion consistently abolished most of the Golgi signal without a corresponding shift to the lysosome, probably because the antigen regions were digested by the lysosome. In contrast, Richards et al. observed striking relocalization of endogenous GNPTαβ to the lysosome in HAP1 cells following TMEM251 deletion, with BafA1 further stabilizing GNPTαβ on the lysosome. However, we and others have independently observed that BafA1 prevents the Golgi exit of GNPTαβ^6,21^. The cause of this discrepancy remains unclear.

### TMEM251 interacts with Golph3 and Retromer to maintain GNPT at the Golgi

How does TMEM251 retain GNPT at the Golgi? Pechincha et al. proposed that GNPT is unstable due to its relatively hydrophilic transmembrane domain. The interaction between TMEM251 and GNPT within the membrane is thought to shield these hydrophilic residues from exposure to the lipid bilayer. Supporting this hypothesis, our alanine scanning results demonstrated that the two transmembrane helices of TMEM251 are critical for its function.

Interestingly, our analysis also revealed that the cytosolic regions adjacent to the transmembrane helices play a significant role in TMEM251 function. Further investigation showed that the N-terminal F^4^XXR^7^ motif of TMEM251 interacts with the COPI adaptor GOLPH3, a key factor in maintaining the TMEM251-GNPT complex at the Golgi. While our paper was under revision, an independent study confirmed that GOLPH3 and its homolog GOLPH3L are crucial for retaining the TMEM251-GNPT complex at the Golgi.

However, additional mechanisms likely contribute to the Golgi retention of TMEM251. Mutating the N-terminal tail to alanine did not completely disrupt the Golgi localization of overexpressed TMEM251 (e.g., the 2-7A mutant, **Fig. 6B**), indicating that Golgi retention is not solely dependent on this region.

Our data also revealed a third layer of complexity to ensure GNPT retention. Specifically, the retromer complex plays a critical role in recycling TMEM251 and GNPT back to the Golgi from post-Golgi compartments. Deleting *VPS35*, a key retromer component, resulted in the mislocalization and degradation of both TMEM251 and GNPT. Importantly, overexpression of TMEM251 partially rescued GNPT stability in *VPS35* KO cells, suggesting that TMEM251 mislocalization is the primary cause of GNPT instability and mislocalization in these cells.

We hypothesize that the C-terminal cytosolic region of TMEM251 may interact with the retromer to mediate the recycling of the TMEM251-GNPT complex. However, demonstrating this interaction has proven challenging. Pull-down assays failed to detect an interaction between WT TMEM251 and the retromer complex, despite VPS35 deletion experiments showing it is critical for TMEM251 localization and GNPT stability. This suggests that the interaction between TMEM251 and the retromer is either transient and weak or mediated by an additional cytosolic adaptor, similar to GOLPH3.

In conclusion, our findings indicate that the trafficking of GNPT within the cell relies on multiple coordinated mechanisms. These include TMEM251-mediated stabilization at the Golgi via transmembrane and cytosolic interactions, GOLPH3-dependent retention, and retromer-mediated retrieval from post-Golgi compartments. Having two distinct methods(i.e., the GNPT-coatomer interaction and the TMEM251-GOLPH3 interaction) for recycling GNPT at the Golgi and a third to retrieve any escaped enzyme ensures the robustness of the M6P pathway, minimizing the risk of its compromise.

## Materials and Methods

### Mammalian cell culture

Cell lines used in this study are listed in Supplemental Table S1. HEK293T (CRL-3216) and HeLa (CCL-2) were purchased from ATCC, and SKMEL-30 (SK1980-526) was purchased from Memorial Sloan Kettering Cancer Center. Cells were cultured in DMEM (Invitrogen) containing 10% Fetal Bovine Serum (Thermo Fisher, and 20% serum for SK-MEL-30 cells), 1% penicillin and streptomycin (Invitrogen), and 1µg/ml plasmocin (Invivogen) at 37°C, 5% CO_2_. All cells were tested negative for mycoplasma using the Mycoalert™ mycoplasma detection kit (Lonza).

### Plasmids

Plasmids used in this study are listed in Supplemental Table S2. The CDS of human TMEM251 was purchased from the DNASU plasmid Repository (Arizona State University).

### Transient transfection

Cells were cultured in DMEM containing 10% (20% for SK-MEL-30 cells) serum media to 40-50% confluency before transfection. Cells were transfected with individual overexpression plasmids (2 µg DNA for a 3.5cm dish) using either Lipofectamine 2000 (Invitrogen) or jetOPTIMUS transfection reagent from Polyplus (Sartorius) according to the manufacturer’s protocol. Cells in 6-well plates were harvested 48 h post-transfection and lysed in 150 µl of buffer A (25 mM Tris-Cl, pH 7.2, 150 mM NaCl, 1% Triton-X 100 and protease inhibitor cocktail).

### Generation of lentiviral stable cell lines

Basically, HEK293T cells were transfected with transfer plasmid, psPAX2 (Addgene 12260), and pMD2.G (Addgene 12259) at a 3.5:3.5:1 ratio using Lipofectamine 2000 according to the manufacturer’s instructions. 72 hours after transfection, the supernatant was collected and passed through a 0.45 µm filter. HEK293T, SK-MEL-30, or HeLa cells were seeded in 6cm dishes and infected with the collected supernatant. After 24 hours of infection, the cells were kept in selection (1 µg/ml for puromycin, and 10 µg/ml for blasticidin) for at least 10 days before subsequent analysis.

### Generation of CRISPR-Cas9 KO and KI cell lines

*TMEM251* and *GNPTAB* knockout cells were generated as described in Ran et al^33^. In brief, sgRNA guides were ligated into pspCas9 (BB)-2A-Puro (Addgene, 48139) or Lenti-multi-CRISPR (Addgene 85402) plasmids. For single colony isolation, cells were transfected with CRISPR-Cas9 knockout plasmids using Lipofectamine 2000 according to the manufacturer’s instruction. 24 hours after transfection, cells were treated with 1µg/ml puromycin (Invitrogen) for 2-3 days. Cells were then diluted into 96-well plates to a final concentration of 0.5 cell per well. The colonies were further screened by western blot analysis, and the candidate clean knockout colonies were verified by sequencing analysis to confirm the indels at their target sites.

The generation of *GNPTAB* knock-in (KI) cell line was described in Zhang et al ^5^. Basically, 300bp homology arms (upstream and downstream from the stop codon) were amplified from the genomic DNA. The 3HA coding sequence was inserted in between the homology arms by overlapping extension. The resulting DNA fragment was ligated into the pGEM-T Easy vector and transfected (4 µg) into HEK293T cells together with CRISPR-Cas9 plasmid (2 µg). 24 hours after transfection, cells were treated with 1µg/ml puromycin (Invitrogen) for 2-3 days. Cells were then diluted into 96-well plates to a final concentration of 0.5 cell per well. The single colonies were screened by PCR using a 3HA internal forward primer and a reverse primer located 600 bp downstream of the stop codon. The candidate KI colonies were further verified by western blot and sequencing analysis.

The following reported sgRNAs were used in this study:

TMEM251 sgRNA1: 5’ – TGTCCACACCCAAAAAGGCA – 3’,

GNPTAB sgRNA1: 5’ – ACTCATTGCGATCTATCGAG – 3’,

GNPTAB sgRNA2 (KI): 5’ – CTTCTATACTCTGATTCGAT – 3’,

### Sample preparation and western blotting

Cells were collected in ice-cold 1X PBS, pelleted at 2700g for 1 min, and lysed in lysis buffer (20 mM Tris pH=8.0, 150 mM NaCl, 1% Triton) containing protease inhibitor cocktail (Bimake) at 4°C for 20 minutes. Cell lysates were centrifuged at 18,000 g for 15 minutes at 4°C. The protein concentration of the supernatant was further measured by Bradford assay (Bio-rad) and normalized. After adding 2X urea sample buffer (150 mM Tris pH 6.8, 6 M Urea, 6% SDS, 40% glycerol, 100 mM DTT, 0.1% Bromophenol blue), samples were heated at 65°C for 8 minutes before further analysis.

Normally, 30 µg of each lysate was loaded and separated on SDS-PAGE gels. Note that for the TMEM251 blot in **Fig. S2**, only 1/20 of the samples were loaded in the TMEM251 overexpression lanes. Protein samples were transferred to a nitrocellulose membrane for western blot analysis. After incubation with primary and secondary antibodies, membranes were scanned using the Odyssey CLx imaging system (LI-COR) or developed with CL-XPosure film (Thermo Scientific).

The following primary antibodies were used for western blotting in this study: rabbit anti-GFP (1:3000, TP401, Torrey Pines Biolabs), mouse anti-actin (1:5000, 66009-1-lg, Proteintech), mouse anti-GAPDH (1:2000, 60004-1-1g, Proteintech), rabbit anti-CTSD (1:1000, 21327-1-AP, Proteintech), rabbit anti-LC3 (1:2000, 14600-1-AP, Proteintech), mouse anti-HA (1:500, 16B12, BioLegend), mouse anti-CTSC (1:500, D-6, Santa Cruz Biotechnology), rabbit anti-LAPTM4A (1:1000, HPA068554-1, Millipore-Sigma), rabbit anti-TMEM251 (1:1000, HPA-48559, Millipore-Sigma), mouse anti-V5 (1:3000, 46-0705, Invitrogen), rabbit anti-Golgin160 (1:1000, 21193-1-AP, Proteintech), Vps35 (1:500, sc-374372, Santa Cruz Biotechnology), Vps26 (1:1000, ab23892, Abcam). Rabbit serum containing antibodies against the α-subunit of GNPTαβ was kindly provided by William Canfield, and the IgG fraction was purified using CaptivA Protein A affinity resin. The generation of single-chain antibodies against M6P was described in Zhang et al. ^5,34^.

The following secondary antibodies were used in this study: goat anti-mouse IRDye 680LT (926-68020), goat anti-mouse IRDye 800CW (926-32210), goat anti-rabbit IRDye 680LT (926-68021), goat anti-rabbit IRDye 800CW (926-32211). These secondary antibodies were purchased from LI-COR Biosciences and used at 1:10,000 dilution.

To detect TMEM251, M6P (scFv M6P), or GNPTAB-3xHA KI, the anti-protein A HRP (PA00-03, Rockland, for TMEM251 and M6P) or mouse HRP (115-035-046, Jackson labs, for GNPTAB-3xHA KI) secondary antibodies were used at 1:10,000 dilution. The signal was detected with the Pierce ECL kit (Thermo Scientific).

### Membrane isolation

The membrane isolation protocol was adapted from Shao and Espenshade^35^, with some modifications. Essentially, cells with 70-80% confluency from a 10cm dish were collected in ice-cold 1X PBS, and pelleted at 2700 g for 1 min. The pelleted cells were resuspended in 1 ml ice-cold membrane isolation buffer (1 mM EDTA and 1 mM EGTA in 1X PBS, with protease inhibitor) and homogenized. The homogenate was centrifuged at 900 g for 5 min at 4°C, and the supernatant was transferred to a new tube and centrifuged at 20,000 g for 20 min at 4 °C to collect membranes. After centrifugation, the membrane pellet was further dissolved in lysis buffer (20 mM Tris pH = 8.0, 150 mM NaCl, and 1% Triton) containing 1X protease inhibitor cocktail (Biomake) at 4°C for 20 minutes. The undissolved membranes were removed by another round of centrifugation at 20,000 g for 15 minutes at 4°C, and the protein concentration from the supernatant was measured by Bradford assay and normalized. Samples were incubated with 2X urea sample buffer samples at 65°C for 8 minutes before western blot analysis.

### Phosphotransferase Activity Assays

The protocol for measuring the GlcNAc-phosphotransferase activity was adapted from van Meel et al.^13^, with some modifications. Essentially, the cells were transfected at 50–60% confluency in six-well plates with a total of 2 μg plasmid DNA (pcDNA3.1 vector, *GNPTAB* cDNA, and *TMEM251* cDNA in the indicated combinations) and 3 μL jetOPTIMUS transfection reagent according to the manufacturer’s protocol. Two days post-transfection, the cells were harvested and lysed in 150 µl of buffer A. Whole cell lysate (50 µg) was assayed in a final volume of 50 µl in buffer containing 50 mM Tris (pH 7.4), 10 mM MgCl_2_, 10 mM MnCl_2_, 2 mg/mL BSA, 2 mM ATP, 75 μM UDP-GlcNAc, and 1 μCi UDP-[^3^H] GlcNAc, with 100 mM α-methylmannoside (α-MM) as acceptor. The reactions were performed at 37 °C for 1 h, quenched with 950 µl of 5 mM EDTA (pH 8.0), and the samples applied to a QAE-Sephadex column (1 ml of packed beads equilibrated with 2 mM Tris, pH 8.0). The columns were washed twice, first with 3 mL of 2 mM Tris (pH 8.0), then with 2 ml of the same buffer. Elution was performed twice, first with 3 ml of 2 mM Tris (pH 8.0) containing 30 mM NaCl, then once again with 2 ml of the same buffer. 8 ml of EcoLite scintillation fluid (MP Biomedicals) was then added to the vials, and the radioactivity in the collected fractions was measured using a Beckman LS6500 scintillation counter. All CPM values that were obtained after subtracting buffer-only background were converted to pmol of GlcNAc-phosphate transferred to α-MM per hour per mg total protein lysate (pmol/hr/mg).

### Lysosomal enzyme assay

Soluble bovine cation-independent mannose 6-phosphate (CI-MPR) was purified from fetal bovine serum as previously described^36^. The purified CI-MPR was conjugated to CNBr-activated Sepharose 4B (Millipore-Sigma) as per the manufacturer’s instructions. For lysosomal enzyme assays, 600 µg of whole cell lysates prepared from either WT HEK293T cells, or from untransfected or 48h transfected TMEM251 knockout, GNPTαβ knockout, or double knockout cells were incubated with equivalent amounts of the CI-MPR affinity beads for 2h at 4 °C, washed twice with cold Tris-buffered saline containing 1%Tx-100 (pH 7.4) (TBST), then assayed for bound lysosomal enzymes as previously described (Gelfman et al., 2007). Briefly, β-hexosaminidase (β-Hex), α-galactosidase (α-Gal) and β-galactosidase (β-Gal) were assayed in a final volume of 50 µl with 5 mM 4-methylumbelliferyl(MU)-N-acetyl-β-d-glucosaminide (Sigma-Aldrich), 5 mM 4-MU-α-d-galactopyranoside and 5 mM 4-MU-β-d-galactopyranoside (Calbiochem, San Diego, CA), respectively, in 50 mM citrate buffer containing 0.5% Triton X-100 (pH 4.5) at 37°C. Reactions were stopped after 1h by addition of 950 µl of 0.4M glycine-NaOH (pH 8.0), samples were vortexed and spun down briefly, and the supernatant transferred to a round-bottomed glass tube. The fluorescence was then measured using a TURNER Model 450 Fluorometer (Barnstead Thermolyne Corporation, Dubuque, Iowa), with excitation and emission wave lengths of 360 and 450 nm, respectively.

### GST pull-down assays

In order to assess the binding of the TMEM251 N-tail to Golph3, GST-Golph3 was first expressed and purified from E. coli BL21 Codon-Plus (RIL) cells (Agilent). For binding assays, 100 µg of the purified GST-Golph3 fusion protein was first immobilized on 50 µl of packed glutathione-agarose beads at room temperature for 1 hour in TBST. The beads were washed once with cold TBST, then 300 µl of whole cell lysates prepared from cells transfected with the various GFP chimeras were added, and the beads tumbled for 2 h at 4 °C. The beads were washed 3 times with cold TBST, and the pellets were resuspended in sample buffer and heated at 100 °C for 10 mins before SDS-gel loading.

### Secretion analysis

Cells were cultured to reach 70-80% confluency to collect secreted proteins from the media. They were washed with serum-free DMEM three times and incubated with serum-free DMEM for 16 hours. The conditioned media were collected and centrifuged at 500 g for 5 min to remove cell debris, filtered with a 0.45 µm filter, and concentrated to ∼100 µl using 10 kDa cutoff Amicon Centrifugal filters (Millipore-Sigma). The protein concentration from the concentrated media was measured by Bradford assay and normalized to its cell lysate. Samples were then mixed with 2 X Urea buffer and heated at 65°C for 8 minutes before further analysis. For the induced secretion assay in **Figure S1B**, the cells were incubated with serum-free DMEM containing 10 mM NH_4_Cl for 16 hours before collection and further processing.

### In-vitro deglycosylation

The deglycosylation assay was performed as described in Venkatarangan et al. ^37^, with some modifications. Basically, for the whole cell lysate, cells were rinsed once with 1x ice-cold PBS and then harvested in 1ml of PBS, split into three equal parts, and centrifuged at 1000 g, 1 min, 4 °C. For deglycosylation assay from the membrane fraction, the pelleted cells were resuspended in 1 ml ice-cold membrane isolation buffer (1 mM EDTA and 1 mM EGTA in 1X PBS, with protease inhibitor) and homogenized. The homogenate was centrifuged at 900 g for 5 min at 4°C, and the supernatant was split into three equal parts, transferred to a new tube, and centrifuged at 20,000 g for 20 min at 4 °C to collect membranes. Two of the three pellets were resuspended in 40μl of lysis buffer (20mM Tris pH=8.0, 150mM NaCl, 1% Triton, 1x PIC), while the third pellet was resuspended in 1X glyco-buffer 2 (50mM sodium phosphate pH 7.5, 1% NP-40, 1X PIC). All pellets were allowed to lyse for 20 minutes at 4°C. Lysates were subsequently centrifuged at 18,000 g, 15 minutes, 4°C, and the protein concentration of the resulting supernatants was measured using Bradford assay. All supernatants were normalized to equal protein concentration. One lysate in the lysis buffer was used as an untreated control. To the second lysate in lysis buffer, 3.9μl of 10X Denaturation buffer (5% SDS, 400mM DTT) was added to 35μl of lysate and boiled at 98°C for 10 minutes. The lysate was then allowed to cool to room temperature for 5 minutes, and a further 4.7μl of 10X glyco-buffer 3 (500mM sodium acetate pH 6) and 2μl of Endo H enzyme were added and incubated at 37°C overnight. To the lysate in glyco-buffer 2, 10μl of glyco-buffer 2 was added to 35μl of the lysate, and 2μl of PNGase F was added further added to it and incubated at 37°C overnight. After incubation, all lysates were mixed with an equal volume of 2 X urea sample buffer, heated at 65°C for 10 minutes before further analysis.

### Fluorescence Protease Protection (FPP) assay

HeLa cells were transfected with GFP tagged construct for 48 hours and grown to a density of about 1-5 × 106 cells/mL (∼80% confluency for a 6 cm dish). The cells were resuspended,nd 10 μL were loaded onto a cell microfluidic plate (Millipore-Sigma, CellASIC ONIX M04S-03) through the cell loading sequence for adherence overnight in 37℃, 5% CO2 incubator^38^. Cells were first washed with KHM buffer (110 mM potassium acetate, 20 mM HEPES, 2 mM MgCl2) for 2 minutes at 6 psi (41.4 kPa), then treated with 20 μM of digitonin for 3 minutes at 6 psi, and with 50 μg/mL proteinase K at 6 psi. Images were captured on the DeltaVision imaging system (GE Healthcare Life Sciences) at intervals of 30 seconds.

### Immunostaining, microscopy, and image processing

Immunostaining was performed as described in Zhang et al.^34^, with some modifications. Cells grown on 1.5 circular glass coverslips were washed with ice-cold 1X PBS and fixed in 4% paraformaldehyde for 10 minutes at room temperature^34^. Cells were permeabilized with 0.3% Triton in PBS for 15 minutes. The samples were blocked in 3% BSA for 30 minutes at room temperature, followed by incubating with primary and secondary antibodies. The cell nucleus was stained using Hoechst (1:8000, Invitrogen). Coverslips were mounted in Fluoromount-G (SouthernBiotech) and cured for 24 hours before imaging.

The following primary antibodies were used for immunostaining in this study: rabbit anti-GNPTAB α subunit (1:100, homemade), rabbit anti-TMEM251 (1:100, HPA-48559, Millipore-Sigma), mouse anti-GM130 (1:200, 610822, BD Biosciences), and rabbit anti-giantin (1:750, 924302, BioLegend).

The following secondary antibodies were used at 1:100: FITC goat anti-rabbit (111-095-003, Jackson ImmunoReseach) and TRITC goat anti-mouse (115-025-003, Jackson ImmunoReseach), sheep anti-mouse (NA931, Cytiva) and donkey anti-rabbit (NA934, Cytiva).

Microscopy was performed with a DeltaVision system (GE Healthcare Life Sciences) or a Dragonfly Confocal Microscope System(Andor Technology) The DeltaVision microscope was equipped with a scientific CMOS camera and an Olympus UPLXAP0100X objective. The following filter sets were used: FITC (excitation, 475/28; emission, 525/48), TRITC (excitation 542/27; emission 594/45), and DAPI (excitation 390/18; emission 435/48). Image acquisition and deconvolution were performed with the SOFTWORX program. For spinning disk confocal microscopy, the following filter sets were used: FITC (excitation, 475/28; emission, 525/48), TRITC (excitation 542/27; emission 594/45), and DAPI (excitation 390/18; emission 435/48). Image acquisition and deconvolution were performed with the Fusion program. Images were further cropped or adjusted using ImageJ (National Institutes of Health).

### Magnetic Isolation of Lysosomes

HEK293T cells were seeded in 150mm dishes containing DMEM with 10% FBS. Once they reached 40% confluency, cells were incubated in DMEM containing 10% FBS, 10 mM HEPES, 10% DexoMAG40 for 24 hours. Following this, cells were incubated in DMEM with 10% FBS for another 24 hours to allow the endocytosis of the magnetic nanoparticles. Following the incubation, cells were rinsed twice and then harvested in 1X ice-cold PBS, using a cell scraper, and spun down at 60xG, 10 minutes, 4°C. The supernatant was discarded, and the pellet was resuspended in 3ml of homogenization buffer(HB: 5mM Tris, 250 mM Sucrose, 1mM EGTA in mass-spectrometry grade water (ThermoFisher), pH=7.4. HB was supplemented with protease inhibitor cocktail (Roche) immediately before use and de-gased). The cell suspension was homogenized in a dounce homogenizer for 30 strokes and then passed through a 22G needle eight times. The homogenate was then centrifuged at 800xG, 10 mins, 4°C. The supernatant was collected, and 80ul was taken as the whole cell lysate (WCL), mixed with equal volume of 2X urea buffer (150mM Tris pH 6.8, 6M Urea, 6% SDS, 40% glycerol, 100mM DTT, 0.1% Bromophenol blue) and heated at 65°C for 10 minutes. The remaining supernatant was allowed to flow freely through the magnetic LS column (130-042-401, Miltenyi Biotech). The column was pre-equilibrated with HB and attached to a platform that provided the magnetic field(130-091-051, Miltenyi Biotech). The column was then washed with 20ml of HB using gravity. The columns were then detached from the magnetic field, and the lysosomes were eluted by flushing 1ml of HB through the column using a syringe. The eluted lysosomes were further concentrated by spinning the eluate at 14,000 rpm, 1 hour, 4°C. The pelleted lysosome fraction was resuspended in 80 ul lysis buffer (20mM Tris pH=8.0, 150mM NaCl, 1% Triton, 1X PIC) and heated at 65°C, for 10 minutes with an equal volume of 2X urea buffer.

### Quantification and statistical analysis

For the quantification of western, the band intensity was quantified using Image Studio software (LI-COR). Graphs were generated using Prism (GraphPad). Statistical analysis was performed with the two-tailed unpaired t-test or one-way ANOVA. Error bars represent the standard deviation. *: ≤0.05, **: ≤0.01, ***: ≤0.001, ****:≤0.0001.

## Data availability

Data supporting this study are provided within the paper and supplementary files. Source data are provided with this paper.

## Acknowledgments

We thank the Li laboratory members for their helpful discussion and technical support. This research is supported by the Protein Folding and Diseases Initiative, a MICHR Pathway Pilot grant, and NIH grants R01HD109346 to M. Li and R01CA008759 to S. Kornfeld.

## Author contributions

Conceptualization: X.Y., B.D., V.V., SK, and M.L.; methodology: X.Y., V.V., B.D., B.J., L.C., and W.Z.; investigation: X.Y.,B.D., V.V., D.H., W.Z., L.C., B.J., L.Y., and B.Z.; writing & editing: V.V., B.D., X.Y., and M.L.; funding acquisition: M.L., and S. K.; resources & supervision: M.L and S.K.; X.Y., BD., and V.V. contributed equally, and all three have the right to list themselves first in bibliographic documents.

## Figure legend

**Figure S1:**
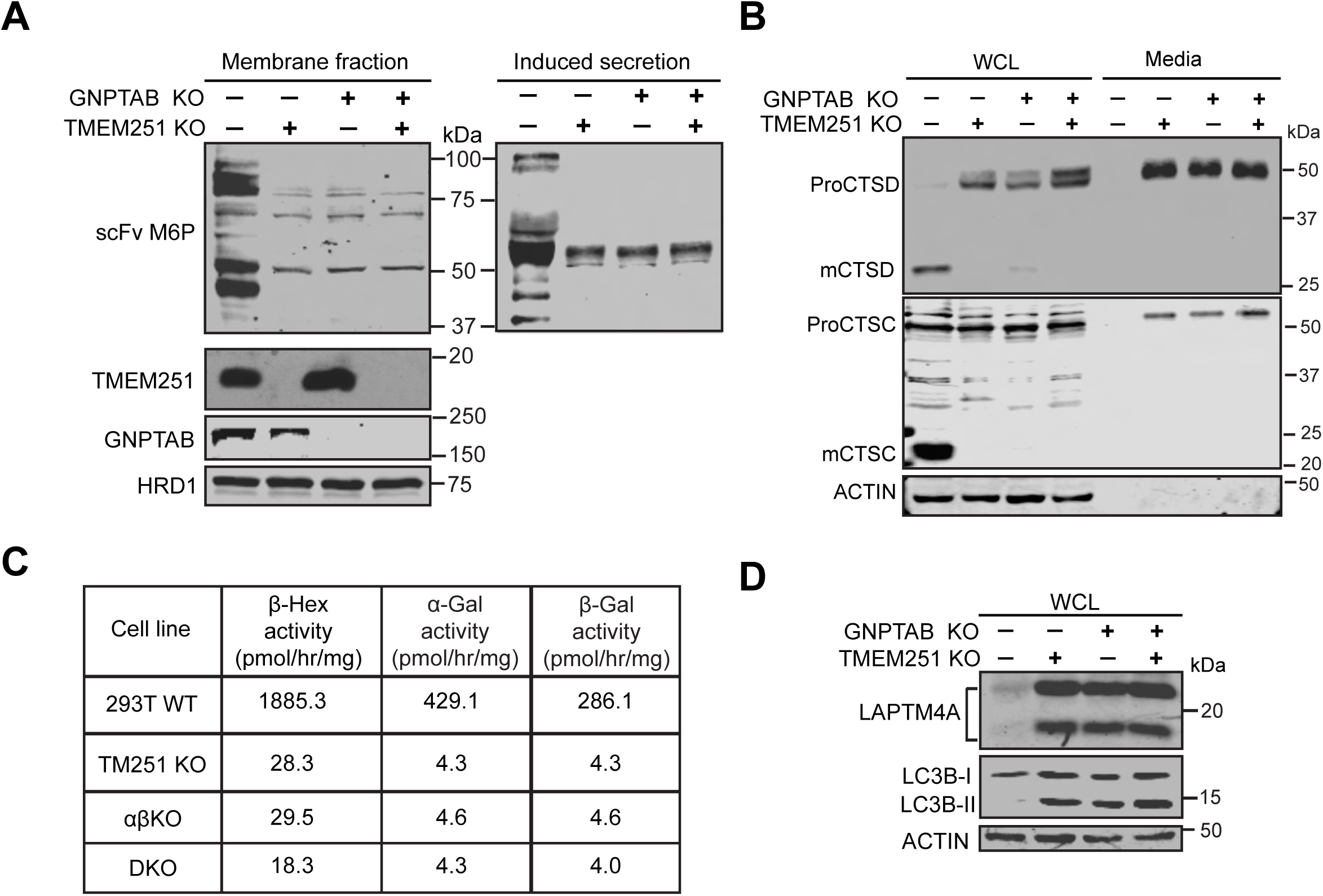
Both GNPTαβ and TMEM251 are essential for the M6P pathway (related to Figure 1) (**A**) Detection of M6P modification in HEK293T WT, TMEM251 KO, GNPTAB KO, and double KO cells using single-chain antibodies against M6P (scFv M6P). (**B**) Immunoblot showing CTSC and CTSD maturation in the whole cell lysate(WCL) and conditioned media of HEK293T WT, *TMEM251* KO, *GNPTAB* KO, and double KO cells. (**C**) A table showing the enzymatic activity of β-Hex, α-gal, and β-gal in HEK293T WT, *TMEM251* KO, *GNPTAB* KO, and double KO cells. Values shown are the average of two assays from two independent transfections. (**D**) Immunoblot showing the accumulation of LAPTM4A and LC3B-II in HEK293T WT, *TMEM251* KO, *GNPTAB* KO, and double KO cells.

**Figure S2:**
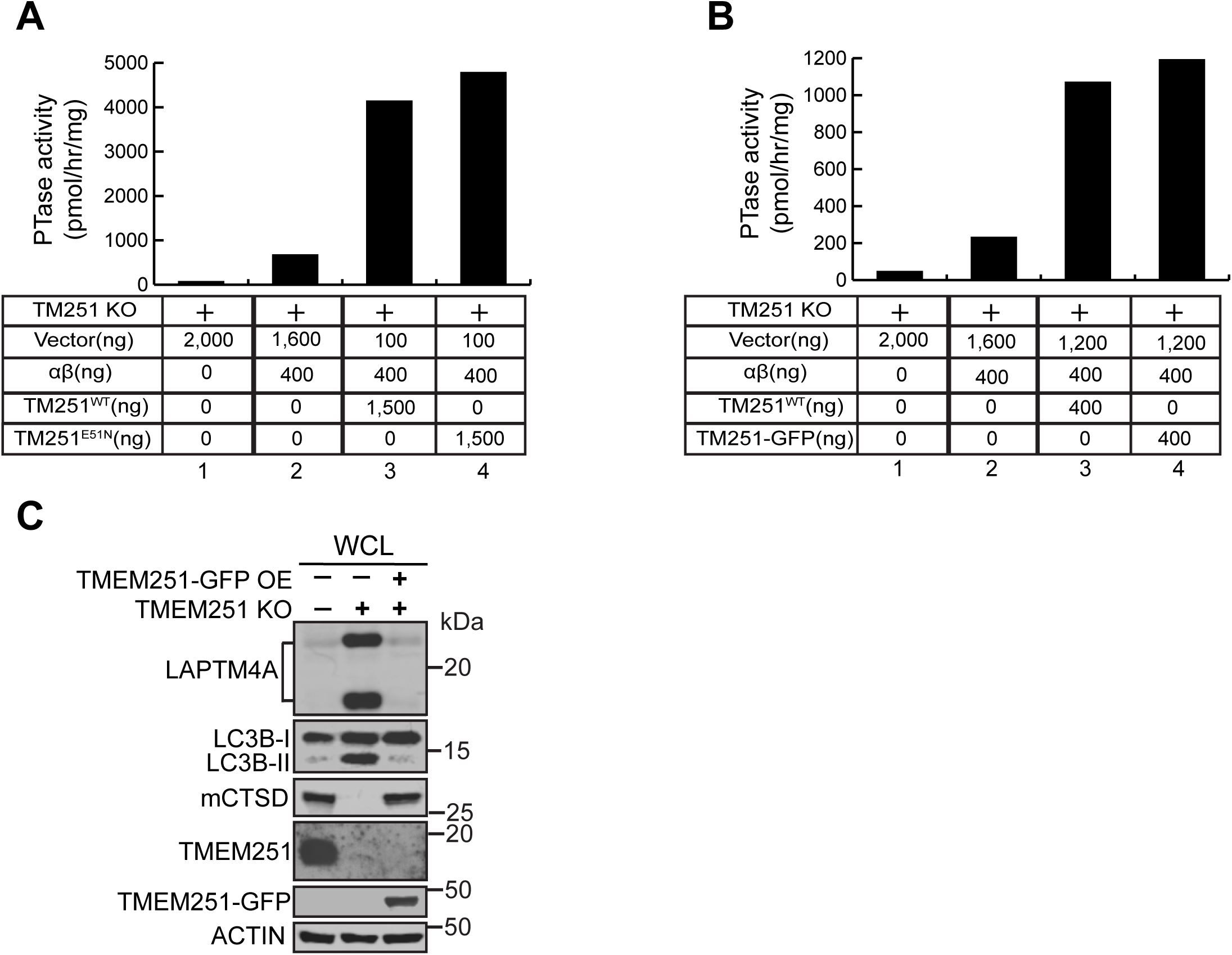
The E51N mutant and TMEM251-GFP are functional (related to Figure 4) (**A**) The E51N mutant restores GNPTαβ phosphotransferase (PTase) activity to levels comparable to WT TMEM251. Values represent the average of two assays from two independent transfections. (**B**) TMEM251-GFP restores GNPTαβ PTase activity to levels similar to WT TMEM251. Values represent the average of two assays from two independent transfections. (**C**) TMEM251-GFP rescues the digestion of lysosomal substrates, including LAPTM4A and LC3-II, in *TMEM251* KO cells.

**Figure S3:**
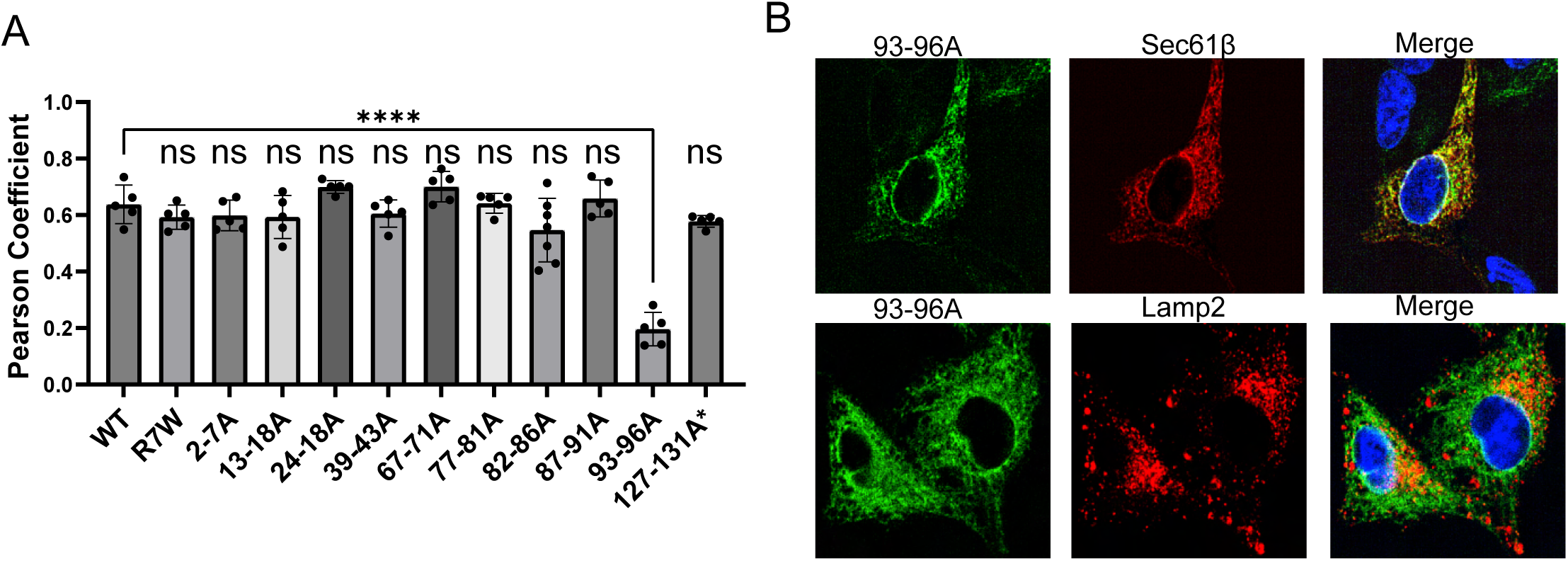
Colocalization analysis of TMEM251 mutants with various organelle markers (related to Figure 5) (**A**) Pearson coefficient analysis showing the colocalization of TMEM251 mutants with the cis-Golgi marker GM130, as shown in Figure 5B. (**B**) The 93-96A TMEM251 mutant colocalizes with the ER marker Sec61β but not the lysosome marker LAMP2.(C)

**Figure S4:**
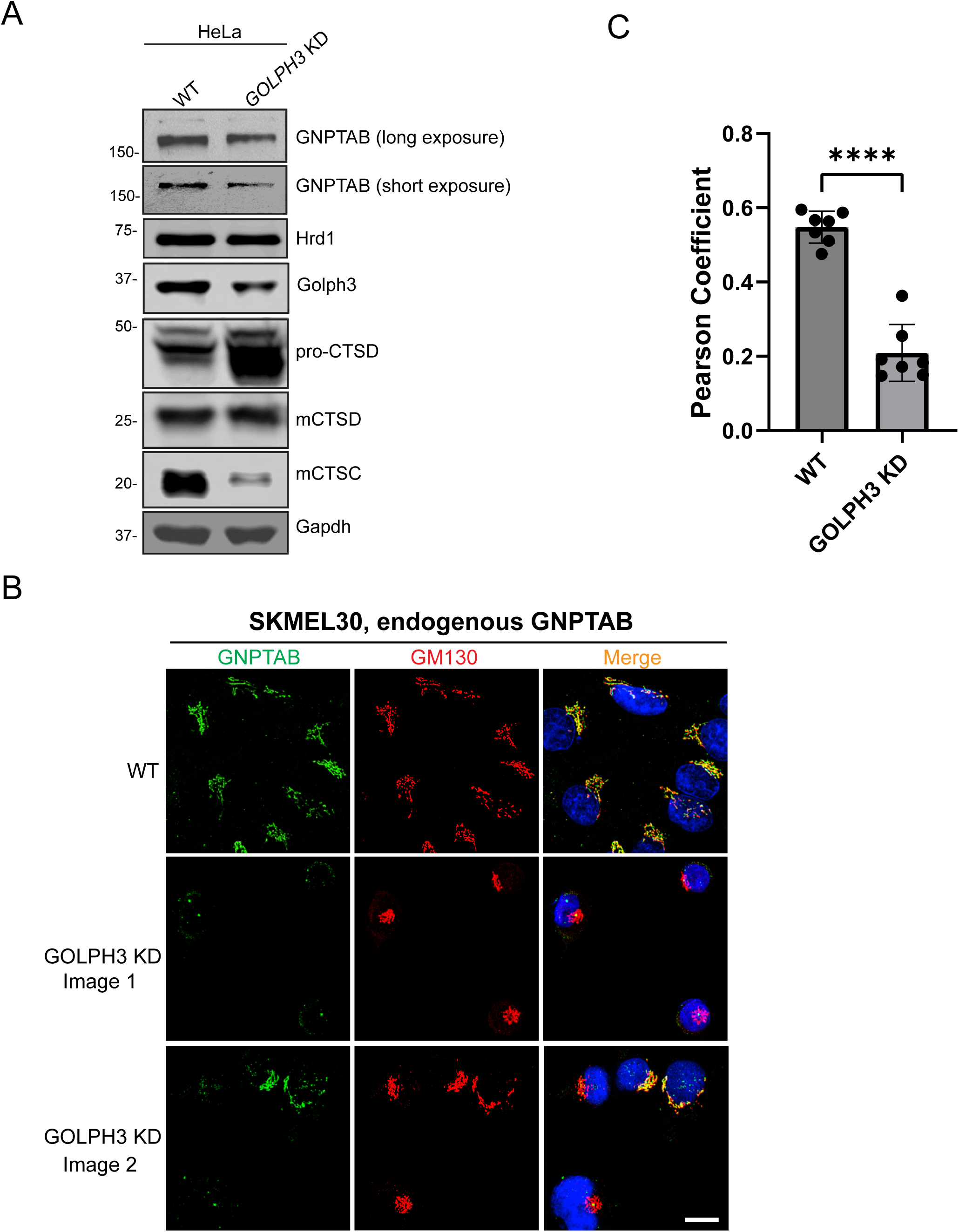
GOLPH3 is critical for the M6P pathway in both HeLa and SKMEL30 cells (related to Figure 6) (**A**) GOLPH3 knockdown in HeLa cells results in reduced endogenous GNPTαβ protein levels and maturation defects in CTSC and CTSD.(**B**) GOLPH3 knockdown in SKMEL30 cells leads to the loss of endogenous GNPTαβ signal at the Golgi.(**C**) Pearson coefficient analysis of (B).

**Supplemental Table 1:**
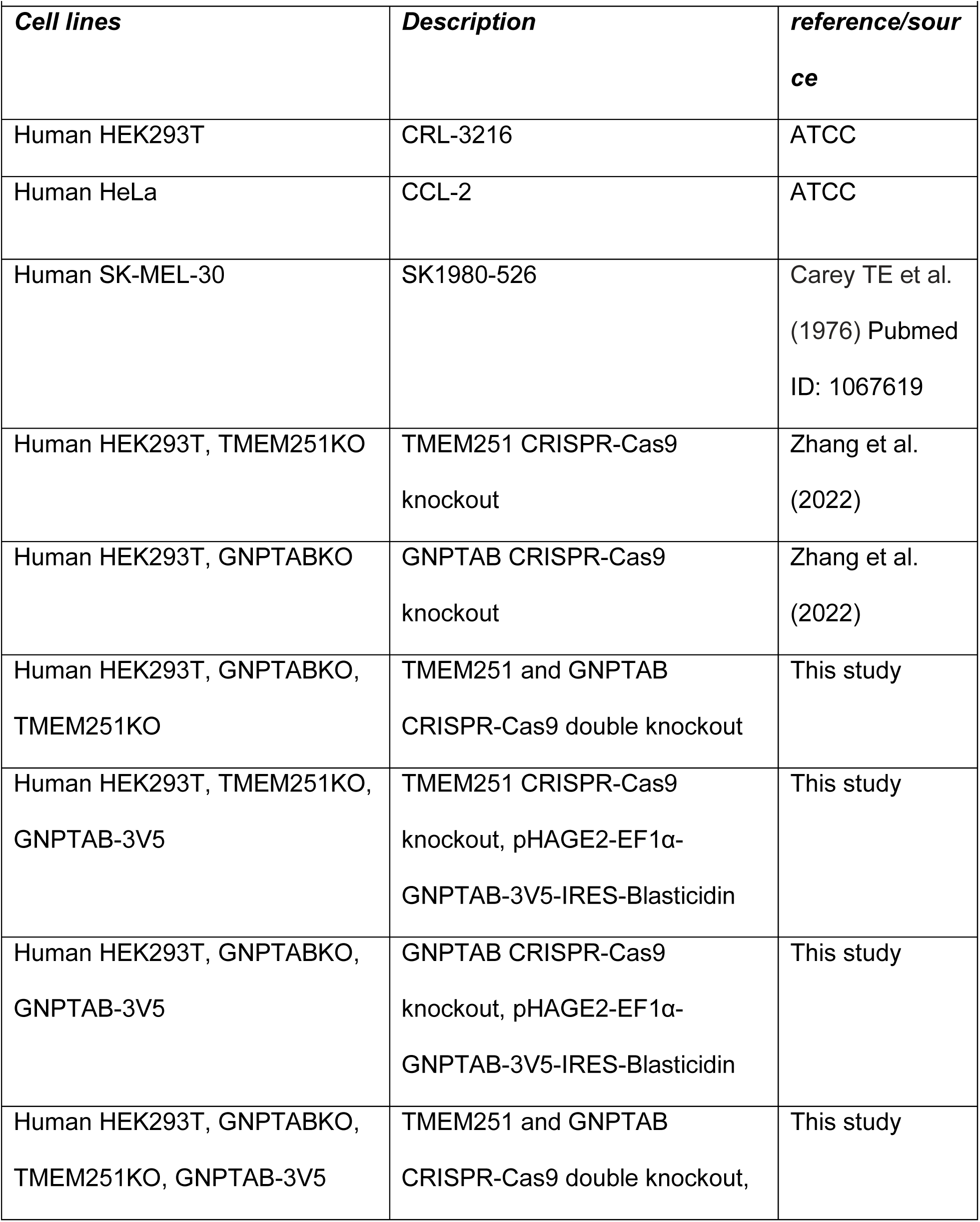
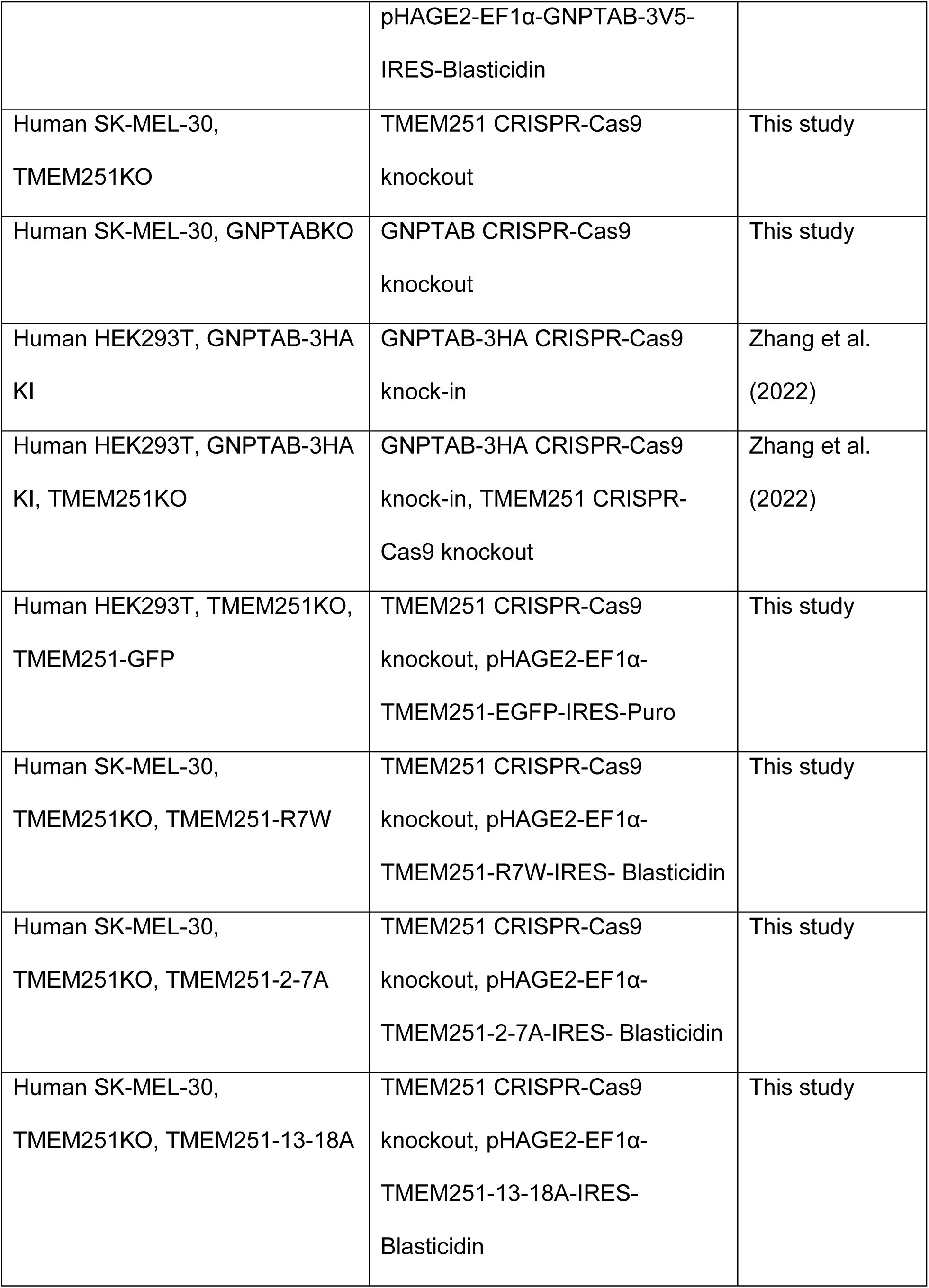
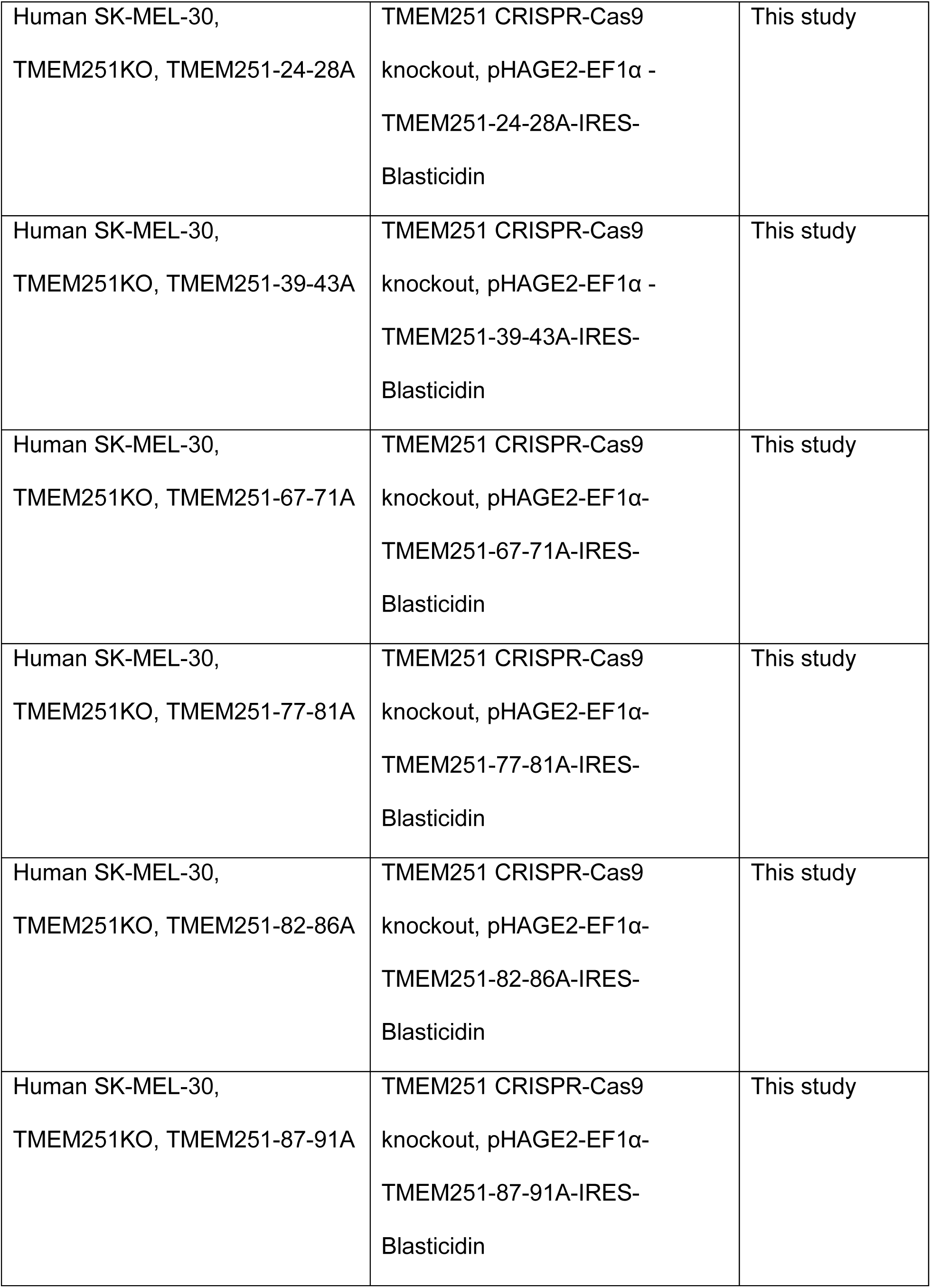
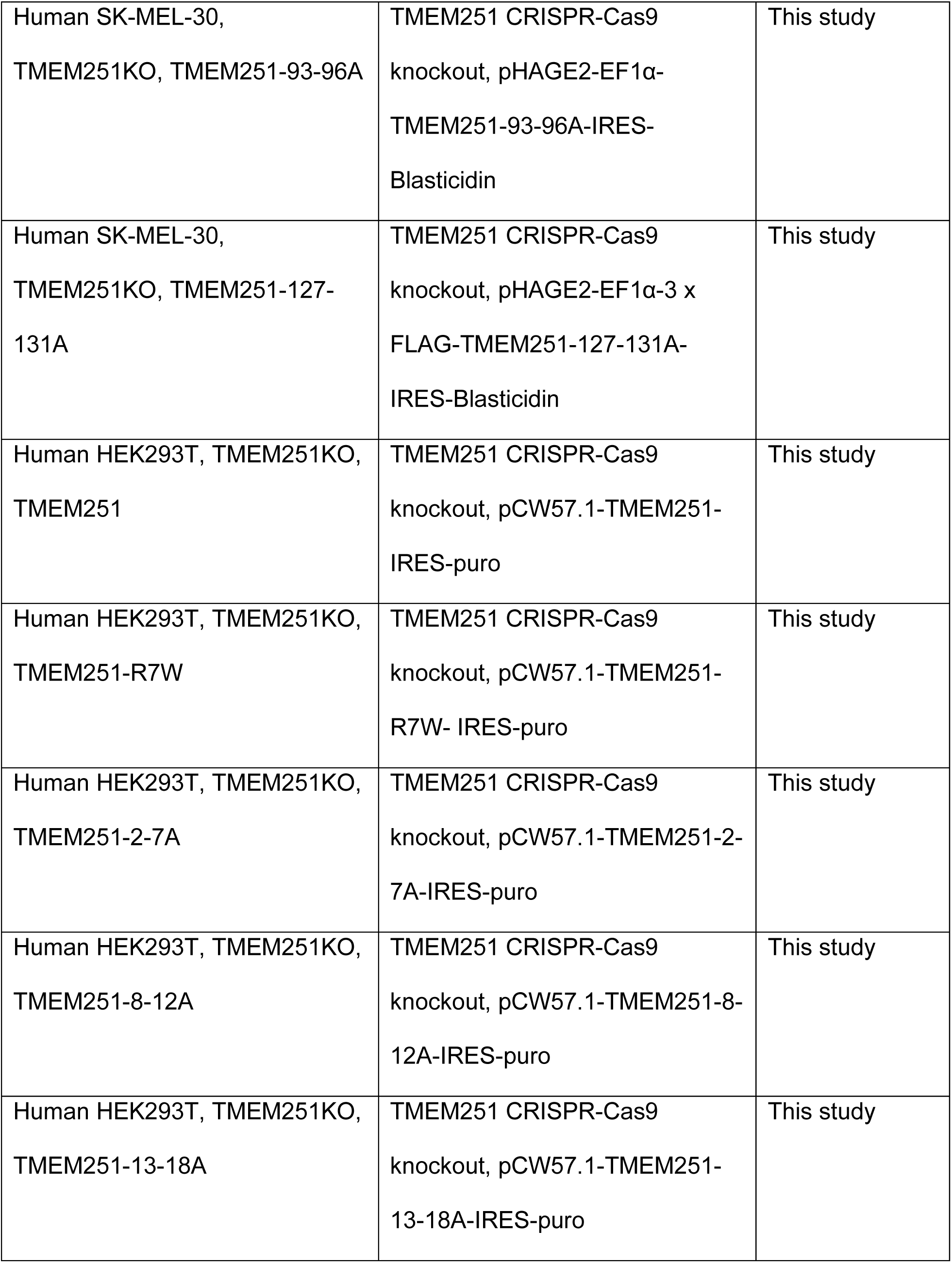
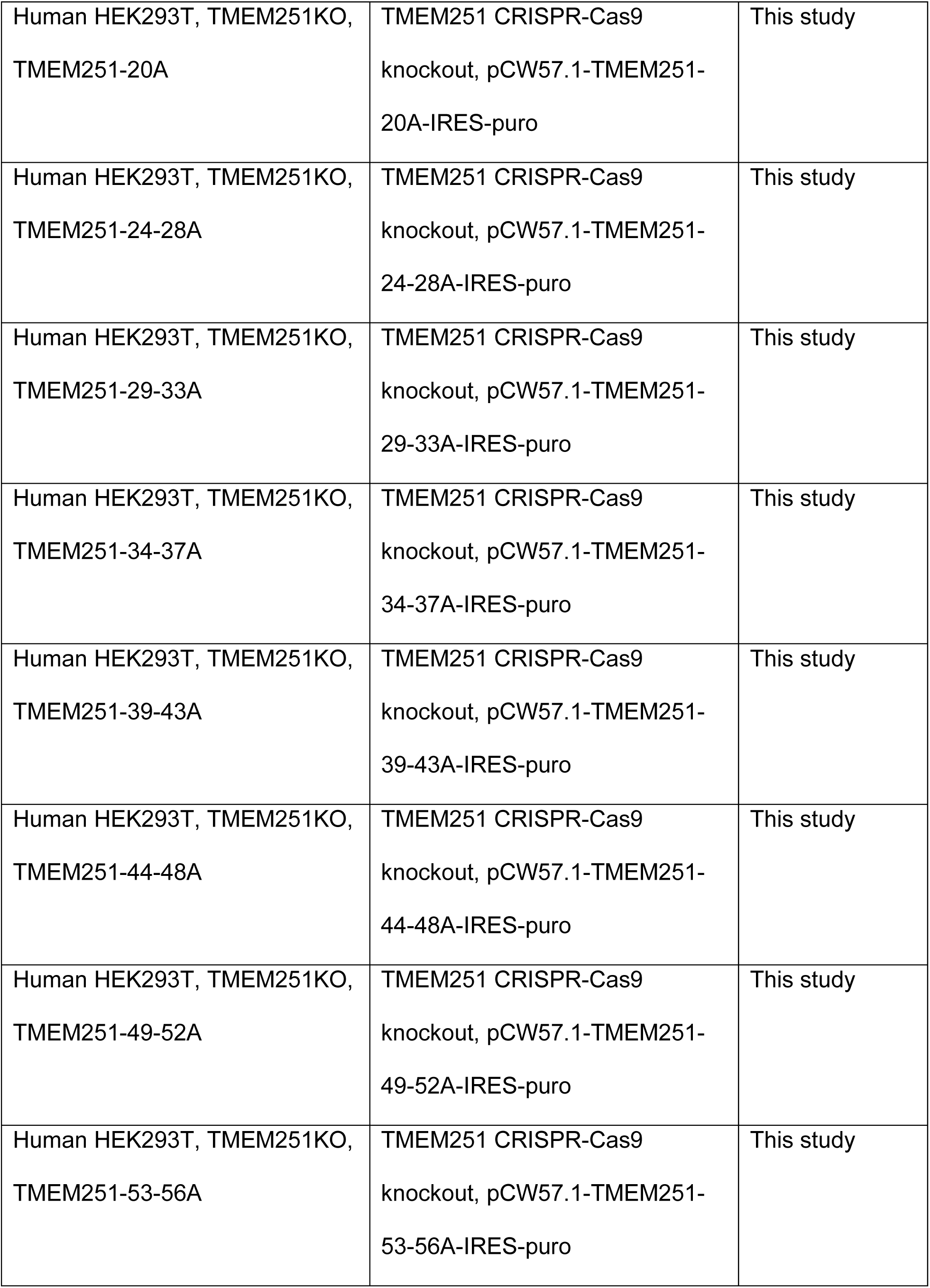
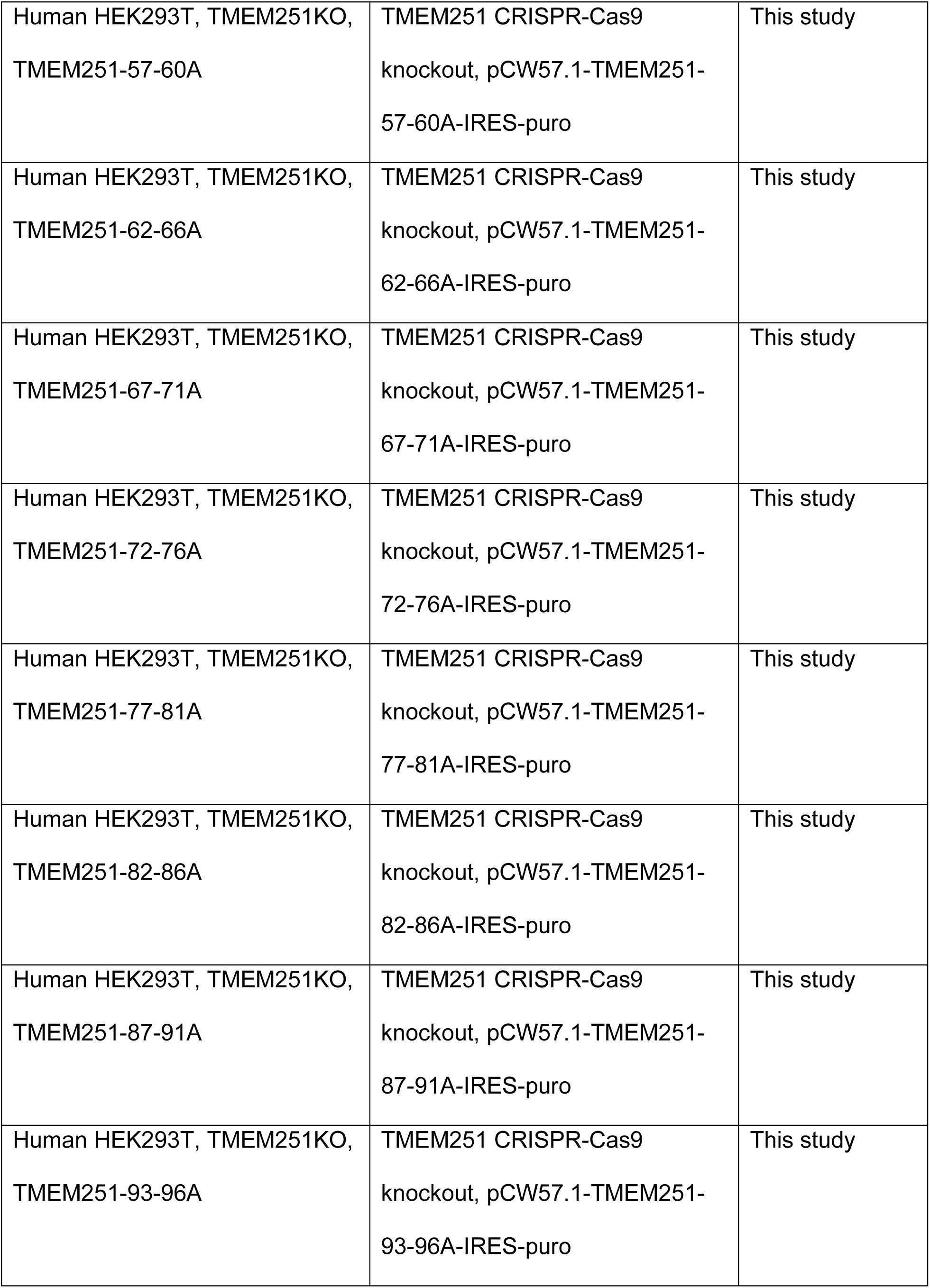
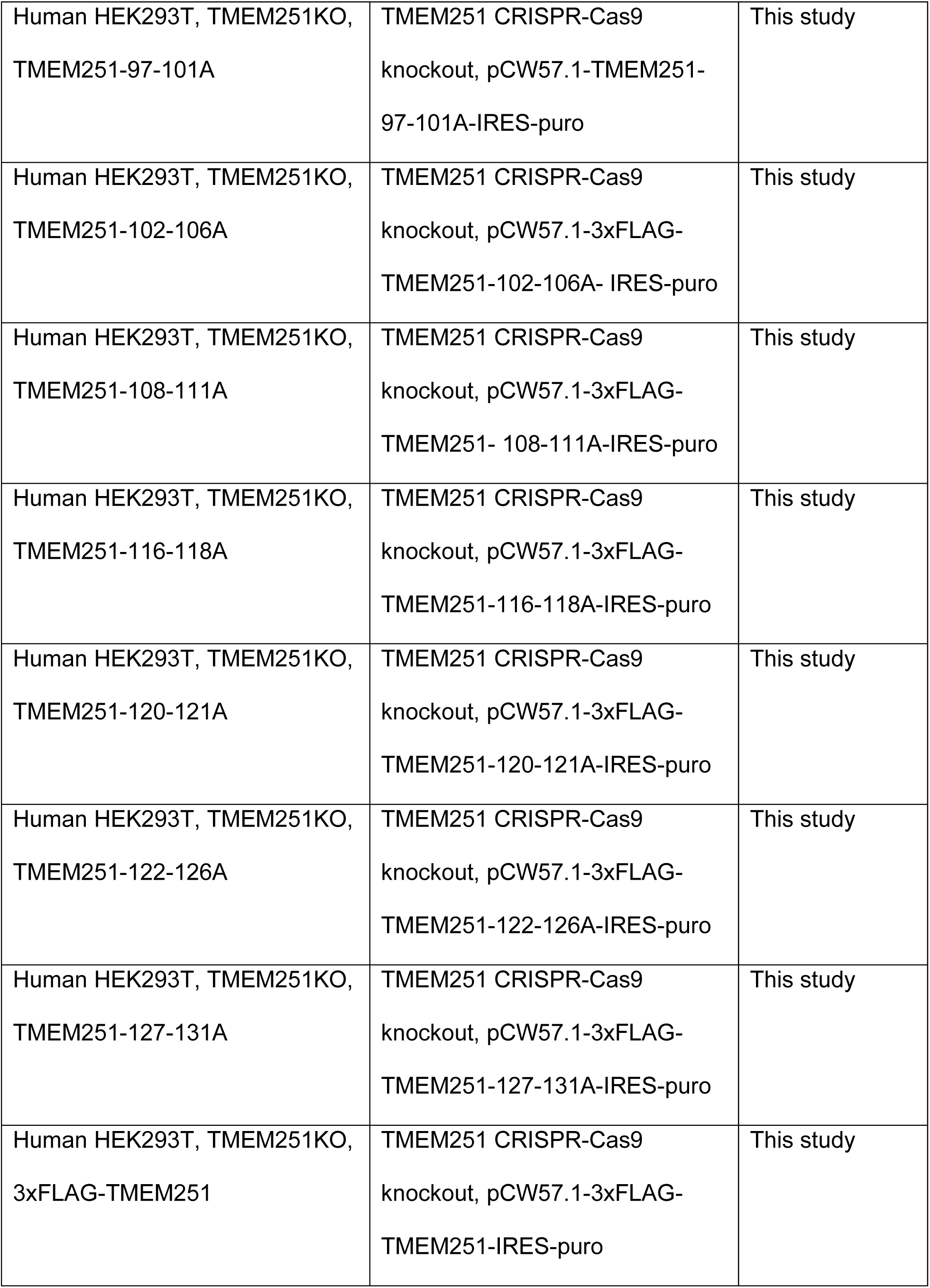
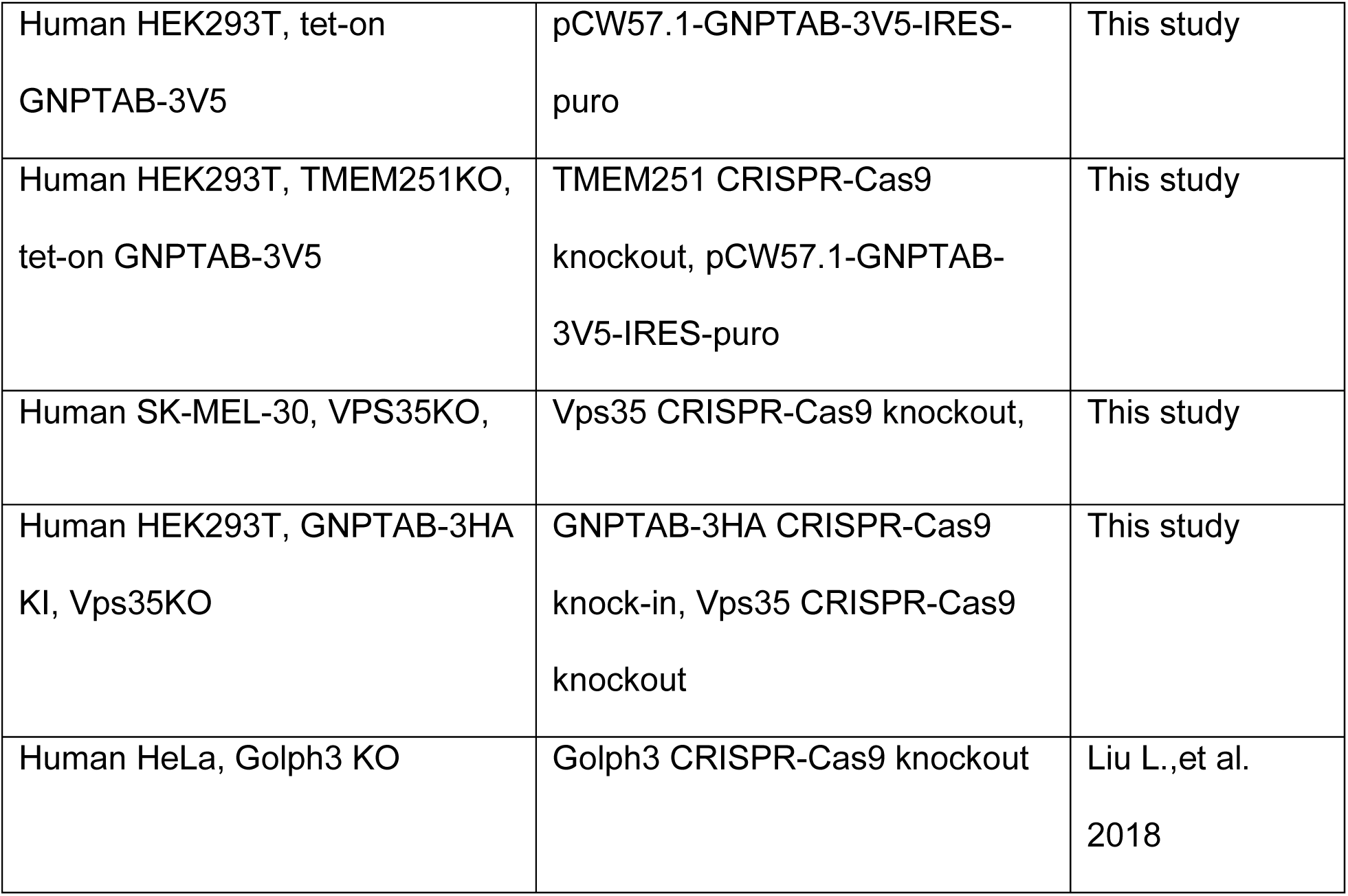
Mammalian cell lines used in this study.

**Supplemental Table 2:** Mammalian plasmids used in this study.

| **Supplemental Table 2: Mammalian plasmids used in this study** |  |  |  |
| --- | --- | --- | --- |
| ***Vector*** | ***Insert*** | ***description*** | ***reference/source*** |
| pSpCas9(BB)-2A-Puro (PX459) |  | CRISPR-Cas9 knockout | Ran et al. 2013<br>Addgene, 48139 |
| psPAX2 |  | Lentiviral packaging<br>plasmid | Addgene 12260 |
| pMD2.G |  | VSV-G envelope | Addgene 12259 |
| Lenti-multi-CRISPR<br>(Mamp194) |  | CRISPR-Cas9 knockout | Cao et al., 2016<br>Addgene 85402 |
| AG949 (Mamp372) | scFv M6P | N-terminal IL-2 signal<br>sequence and C-terminal<br>Fc region of Rabbit IgG | University of<br>Geneva,<br>Zhang et al., 2022 |
| Man1A1-GFP | Man1A1 |  |  |
| pEGFP-N1 | CTNS | CMV promoter, C-terminal GFP | This study |
| pEGFP-N1 | TMEM251 | CMV promoter, C-terminal GFP | This study |
| pcDNA3.1 | GNPTAB-V5-6His | CMV promoter | This study |
| pcDNA3.1 | TMEM251 | CMV promoter | This study |
| pcDNA3.1 | TMEM251-E41N | CMV promoter | This study |

## References

1 Ghosh, P., Dahms, N. M. & Kornfeld, S. Mannose 6-phosphate receptors: new twists in the tale. Nat Rev Mol Cell Biol 4, 202–212, doi:10.1038/nrm1050 (2003).

2 Settembre, C. & Perera, R. M. Lysosomes as coordinators of cellular catabolism, metabolic signalling and organ physiology. Nat Rev Mol Cell Biol, doi:10.1038/s41580-023-00676-x (2023).

3 Khan, S. A. & Tomatsu, S. C. Mucolipidoses Overview: Past, Present, and Future. Int J Mol Sci 21, doi:10.3390/ijms21186812 (2020).

4 Zhang, B., Yang, X. & Li, M. LYSET/TMEM251/GCAF is critical for autophagy and lysosomal function by regulating the mannose-6-phosphate (M6P) pathway. Autophagy, 1–3, doi:10.1080/15548627.2023.2167375 (2023).

5 Zhang, W. et al. GCAF(TMEM251) regulates lysosome biogenesis by activating the mannose-6-phosphate pathway. Nat Commun 13, 5351, doi:10.1038/s41467-022-33025-1 (2022).

6 Pechincha, C. et al. Lysosomal enzyme trafficking factor LYSET enables nutritional usage of extracellular proteins. Science 378, eabn5637, doi:10.1126/science.abn5637 (2022).

7 Richards, C. M. et al. The human disease gene LYSET is essential for lysosomal enzyme transport and viral infection. Science 378, eabn5648, doi:10.1126/science.abn5648 (2022).

8 Ain, N. U. et al. Biallelic TMEM251 variants in patients with severe skeletal dysplasia and extreme short stature. Hum Mutat 42, 89–101, doi:10.1002/humu.24139 (2021).

9 Braulke, T., Carette, J. E. & Palm, W. Lysosomal enzyme trafficking: from molecular mechanisms to human diseases. Trends Cell Biol, doi:10.1016/j.tcb.2023.06.005 (2023).

10 Qiao, W., Richards, C. M. & Jabs, S. LYSET/TMEM251-a novel key component of the mannose 6-phosphate pathway. Autophagy 19, 2143–2145, doi:10.1080/15548627.2023.2167376 (2023).

11 van Meel, E., Qian, Y. & Kornfeld, S. A. Mislocalization of phosphotransferase as a cause of mucolipidosis III alphabeta. Proc Natl Acad Sci U S A 111, 3532–3537, doi:10.1073/pnas.1401417111 (2014).

12 Muller-Loennies, S., Galliciotti, G., Kollmann, K., Glatzel, M. & Braulke, T. A novel single-chain antibody fragment for detection of mannose 6-phosphate-containing proteins: application in mucolipidosis type II patients and mice. Am J Pathol 177, 240–247, doi:10.2353/ajpath.2010.090954 (2010).

13 van Meel, E. et al. Multiple Domains of GlcNAc-1-phosphotransferase Mediate Recognition of Lysosomal Enzymes. J Biol Chem 291, 8295–8307, doi:10.1074/jbc.M116.714568 (2016).

14 Li, H. et al. Structure of the human GlcNAc-1-phosphotransferase alphabeta subunits reveals regulatory mechanism for lysosomal enzyme glycan phosphorylation. Nat Struct Mol Biol, doi:10.1038/s41594-022-00748-0 (2022).

15 Thomas, G. Furin at the cutting edge: from protein traffic to embryogenesis and disease. Nat Rev Mol Cell Biol 3, 753–766, doi:10.1038/nrm934 (2002).

16 Kisselev, A. F., Akopian, T. N. & Goldberg, A. L. Range of sizes of peptide products generated during degradation of different proteins by archaeal proteasomes. J Biol Chem 273, 1982–1989, doi:10.1074/jbc.273.4.1982 (1998).

17 Kisselev, A. F., Akopian, T. N., Woo, K. M. & Goldberg, A. L. The sizes of peptides generated from protein by mammalian 26 and 20 S proteasomes. Implications for understanding the degradative mechanism and antigen presentation. J Biol Chem 274, 3363–3371 (1999).

18 Jumper, J. et al. Highly accurate protein structure prediction with AlphaFold. Nature 596, 583–589, doi:10.1038/s41586-021-03819-2 (2021).

19 Tsirigos, K. D., Peters, C., Shu, N., Kall, L. & Elofsson, A. The TOPCONS web server for consensus prediction of membrane protein topology and signal peptides. Nucleic Acids Res 43, W401–407, doi:10.1093/nar/gkv485 (2015).

20 Viklund, H. & Elofsson, A. OCTOPUS: improving topology prediction by two-track ANN-based preference scores and an extended topological grammar. Bioinformatics 24, 1662–1668, doi:10.1093/bioinformatics/btn221 (2008).

21 Liu, L., Doray, B. & Kornfeld, S. Recycling of Golgi glycosyltransferases requires direct binding to coatomer. Proc Natl Acad Sci U S A 115, 8984–8989, doi:10.1073/pnas.1810291115 (2018).

22 Welch, L. G., Peak-Chew, S. Y., Begum, F., Stevens, T. J. & Munro, S. GOLPH3 and GOLPH3L are broad-spectrum COPI adaptors for sorting into intra-Golgi transport vesicles. J Cell Biol 220, doi:10.1083/jcb.202106115 (2021).

23 Ali, M. F., Chachadi, V. B., Petrosyan, A. & Cheng, P. W. Golgi phosphoprotein 3 determines cell binding properties under dynamic flow by controlling Golgi localization of core 2 N-acetylglucosaminyltransferase 1. J Biol Chem 287, 39564–39577, doi:10.1074/jbc.M112.346528 (2012).

24 Tu, L., Tai, W. C., Chen, L. & Banfield, D. K. Signal-mediated dynamic retention of glycosyltransferases in the Golgi. Science 321, 404–407, doi:10.1126/science.1159411 (2008).

25 Rizzo, R. et al. Golgi maturation-dependent glycoenzyme recycling controls glycosphingolipid biosynthesis and cell growth via GOLPH3. EMBO J 40, e107238, doi:10.15252/embj.2020107238 (2021).

26 Seaman, M. N., Burd, C. G. & Emr, S. D. Receptor signalling and the regulation of endocytic membrane transport. Curr Opin Cell Biol 8, 549–556 (1996).

27 Seaman, M. N., Marcusson, E. G., Cereghino, J. L. & Emr, S. D. Endosome to Golgi retrieval of the vacuolar protein sorting receptor, Vps10p, requires the function of the VPS29, VPS30, and VPS35 gene products. J Cell Biol 137, 79–92 (1997).

28 Suzuki, S. W., Chuang, Y. S., Li, M., Seaman, M. N. J. & Emr, S. D. A bipartite sorting signal ensures specificity of retromer complex in membrane protein recycling. J Cell Biol 218, 2876–2886, doi:10.1083/jcb.201901019 (2019).

29 Seaman, M. N. The retromer complex - endosomal protein recycling and beyond. J Cell Sci 125, 4693–4702, doi:10.1242/jcs.103440 (2012).

30 Arighi, C. N., Hartnell, L. M., Aguilar, R. C., Haft, C. R. & Bonifacino, J. S. Role of the mammalian retromer in sorting of the cation-independent mannose 6-phosphate receptor. J Cell Biol 165, 123–133, doi:10.1083/jcb.200312055 (2004).

31 Gorelik, A., Illes, K., Bui, K. H. & Nagar, B. Structures of the mannose-6-phosphate pathway enzyme, GlcNAc-1-phosphotransferase. Proc Natl Acad Sci U S A 119, e2203518119, doi:10.1073/pnas.2203518119 (2022).

32 Du, S. et al. Structural insights into how GlcNAc-1-phosphotransferase directs lysosomal protein transport. J Biol Chem 298, 101702, doi:10.1016/j.jbc.2022.101702 (2022).

33 Ran, F. A. et al. Genome engineering using the CRISPR-Cas9 system. Nat Protoc 8, 2281–2308, doi:10.1038/nprot.2013.143 (2013).

34 Zhang, W. et al. A conserved ubiquitin- and ESCRT-dependent pathway internalizes human lysosomal membrane proteins for degradation. PLoS Biol 19, e3001361, doi:10.1371/journal.pbio.3001361 (2021).

35 Shao, W. & Espenshade, P. J. Sterol regulatory element-binding protein (SREBP) cleavage regulates Golgi-to-endoplasmic reticulum recycling of SREBP cleavage-activating protein (SCAP). J Biol Chem 289, 7547–7557, doi:10.1074/jbc.M113.545699 (2014).

36 Valenzano, K. J., Remmler, J. & Lobel, P. Soluble insulin-like growth factor II/mannose 6-phosphate receptor carries multiple high molecular weight forms of insulin-like growth factor II in fetal bovine serum. J Biol Chem 270, 16441–16448, doi:10.1074/jbc.270.27.16441 (1995).

37 Venkatarangan, V. et al. ER-associated degradation in cystinosis pathogenesis and the prospects of precision medicine. J Clin Invest 133, doi:10.1172/JCI169551 (2023).

38 Yang, X. et al. ESCRT, not intralumenal fragments, sorts ubiquitinated vacuole membrane proteins for degradation. J Cell Biol 220, doi:10.1083/jcb.202012104 (2021).

39 Robinson, J. S., Klionsky, D. J., Banta, L. M. & Emr, S. D. Protein sorting in Saccharomyces cerevisiae: isolation of mutants defective in the delivery and processing of multiple vacuolar hydrolases. Mol Cell Biol 8, 4936–4948 (1988).

